# Chromosomal Instability in Human Trophoblast Stem Cells and Placentas

**DOI:** 10.1101/2024.11.15.623799

**Authors:** Danyang Wang, Andrew Cearlock, Katherine Lane, Ian Jan, Rajiv McCoy, Min Yang

**Affiliations:** Department of Obstetrics & Gynecology, University of Washington, Seattle, WA, USA; Institute for Stem Cell & Regenerative Medicine, University of Washington, Seattle, WA, USA; Department of Bioengineering, University of Washington, Seattle, WA, USA; Department of Biology, Johns Hopkins University, Baltimore, MD, USA; Brotman Baty Institute for Precision Medicine, Seattle, WA, USA; Washington National Primate Center, Seattle, WA, USA

## Abstract

The human placenta, a unique tumor-like organ, is typically thought to exhibit rare aneuploidy associated with adverse pregnancy outcomes. Discrepancies in reported aneuploidy prevalence in placenta likely stem from limitations in modeling and the resolution of detection methods. Here, we used isogenic trophoblast stem cells (TSCs) derived from both naïve and primed human pluripotent stem cells (hPSCs) to reveal the spontaneous occurrence of aneuploidy, suggesting chromosomal instability (CIN) as an inherent feature of the trophoblast lineage. We identified potential pathways contributing to the occurrence and tolerance of CIN. These findings were further validated using single cell multiome data from human placentas, where we observed a high prevalence of heterogeneous aneuploidy across trophoblast cells. Despite extensive chromosomal abnormalities, TSCs maintained their proliferative and differentiation capacities, suggesting that CIN is a typical aspect of placental development. Our study challenges the traditional view of aneuploidy in the placenta and provides new insights into the role of CIN in normal placental function.

## Introduction

Placental aneuploidy has been considered a detrimental factor associated with adverse pregnancy outcomes and placental dysfunction^1^. Chromosomal aberrations confined to the placenta are found in approximately 1–2% of pregnancies, and these abnormalities, however, do not always impact fetal development^2–5^.

The human placenta, a critical organ for fetal growth and development in utero, is predominantly composed of trophoblasts (TBs), which originate from the trophectoderm (TE). Cytotrophoblasts (CTs), a specific type of trophoblast, play a pivotal role in embedding the embryo into the uterine lining, marking the onset of placental formation. CTs proliferate to establish the foundational structures of the placenta and differentiate into two key cell types: syncytiotrophoblasts (STs), which form the outer layer of the placental villi responsible for nutrient and gas exchange, and extravillous trophoblasts (EVTs), which invade the maternal uterine wall to anchor the placenta and facilitate the establishment of maternal-fetal circulation.

Studies by Weier et al. identified a high level of numerical chromosomal abnormalities in a subtype of cytotrophoblasts from pregnancies terminated for non-medical reasons and, more recently, revealed extensive chromosomal abnormalities in invasive CTs, suggesting that such heterogeneity may confer a selective advantage for invasion^6,7^. However, comprehensive data to support the bold hypothesis that aneuploidy may be a common feature of placental development are lacking, as most current evidence primarily detected aneuploidy in abnormal pregnancies, such as those involving intrauterine growth restriction or preeclampsia^8,9^. This discrepancy in data likely arises from limited research on the placenta and the absence of robust single-cell resolution analyses. Given the dynamic and error-prone nature of human embryogenesis, understanding what constitutes “normal” and acceptable variation in the placenta is crucial for diagnostic purposes. This raises the fundamental question: what is “normal” for the placenta? Surprisingly, despite its critical role in pregnancy, the cellular phenotypes of placental cells remain largely unexplored, posing a significant barrier to improving clinical pregnancy prognostics.

Recent advancements have enabled the in vitro derivation of CTs as trophoblast stem cells (TSCs) from both primary tissues^10,11^ and human pluripotent stem cells (hPSCs)^12–25^. These TSCs have demonstrated the capacity to differentiate into various trophoblast subtypes and form organoids, providing a novel platform for studying early placental development.

To explore the cellular phenotype of aneuploidy in placental lineages, we established isogenic TSC lines using multiple methodologies from both primed and naïve hPSCs, as well as from preimplantation embryo model blastoids. Intriguingly, we observed a high frequency of spontaneous aneuploidy across all TSC lines, regardless of derivation method, suggesting chromosomal instability (CIN) as an intrinsic trait of the trophoblast lineage. Key pathways potentially driving this phenomenon were identified. Similarly, first-trimester and term placental tissues showed accumulated chromosomal abnormalities. These findings may reshape our understanding of normal placental cellular physiology and open new avenues for investigating the implications of CIN in placental development.

## Results

### Trophoblast Stem Cells Can Be Derived from Isogenic Primed, Naïve hPSCs, and Blastoids

We adopted and optimized several approaches for generating TSCs from both naïve and primed hPSCs. To minimize cell line-specific bias, we used the same isogenic parental hPSCs (RUES2, NIHhESC-09-0013). As no direct comparisons of these methods have been previously made, our isogenic data provide a foundational reference.

All trophoblast lineages arise from the blastocyst’s TE. When deriving TSCs from naïve hPSCs, some approaches require an intermediate TE induction phase^17^ or sorting to obtain a homogenous TSC population^14^. However, homogeneous TSCs have also been reported to be derived directly from naïve hPSCs without a TE intermediate or sorting^13^.Unlike primed hPSCs, naïve hPSCs cannot undergo BMP4-directed trophoblast differentiation without capacitation or re-priming^13^. Instead, naïve hPSCs respond robustly to induction with TGF-beta pathway inhibitor A83 and ERK inhibitor PD0325901^17^. This BMP4 independence in trophoblast specification of naïve hPSCs aligns with BMP4’s lack of effect on preimplantation embryos^26^. We reprogrammed primed hPSCs to a stabilized naïve state in PXGL medium^27^ (Figure 1A). To generate naïve TSCs (nTSC), naïve hPSCs were then transitioned to TSC medium on MEFs, moving to feeder-free culture by the second passage. This process generated a uniform, highly proliferative population resembling primary TSCs morphologically^10^, expressing key TSC markers GATA3, KRT7, GATA2, and TFAP2A at protein and transcript levels (Figures 1B, 1C, and S1B).

**Figure 1.**
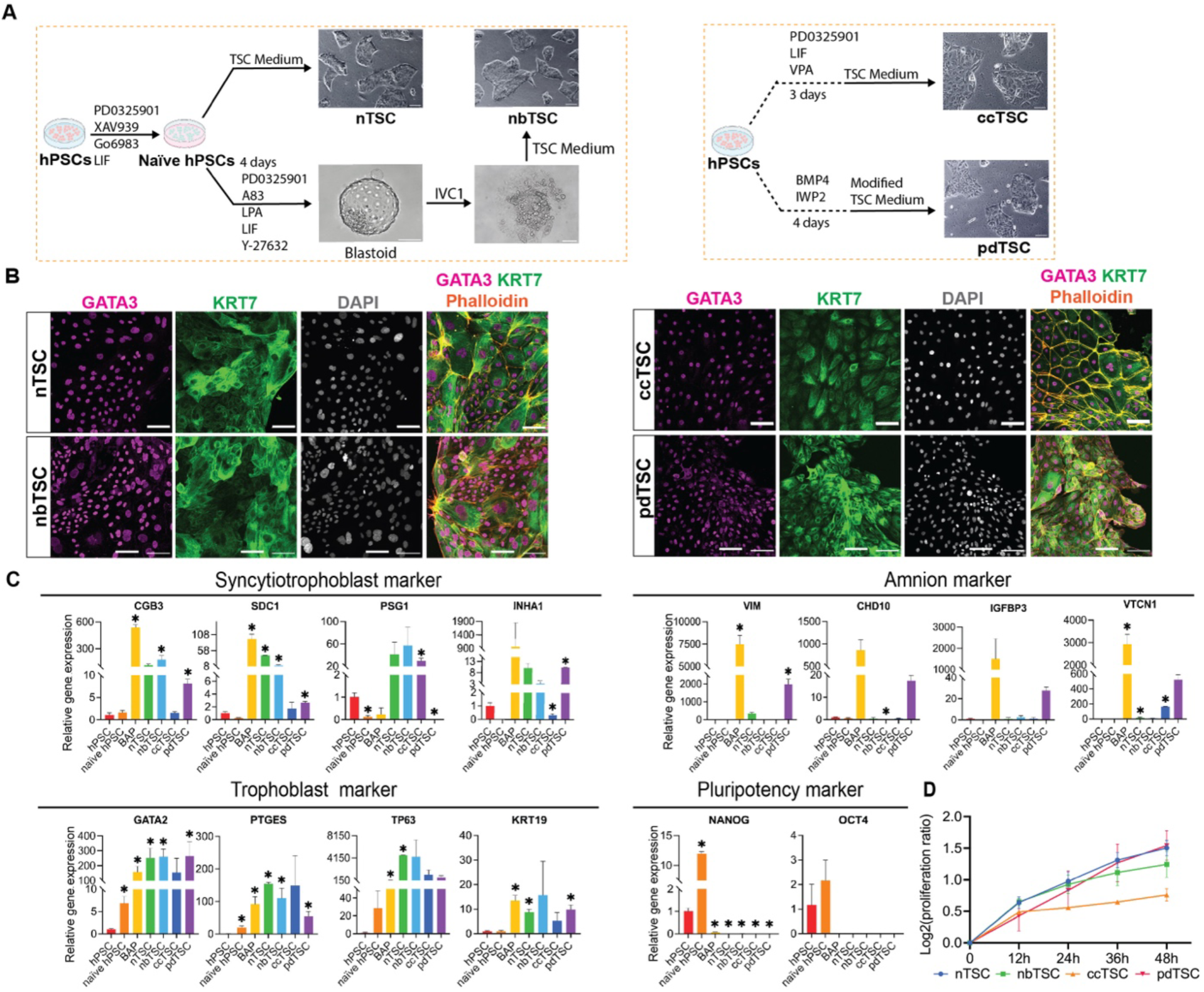
Derivation of TSCs from naïve and primed hPSCs. (A) Schematic representation of protocols for deriving four types of TSCs. nTSC: directly derived from naïve hPSCs; nbTSC: derived from attached naïve hPSC-derived blastoids; ccTSC: generated from primed hPSCs via chemical conversion with HDACi VPA, PD0325901, and LIF, involving an intermediate state of epigenetic reprogramming; pdTSC: induced from primed hPSCs with initial BMP4 and WNTi (IWP2) treatment to prime cells toward the trophectoderm phase. (B) Immunostaining of GATA3 and KRT7 in nTSCs, nbTSCs, ccTSCs, and pdTSCs. Phalloidin and DAPI staining for cytoskeleton and nucleus. Scale bar, 100 μm. (C) Quantitative gene expression analysis of pluripotency, syncytiotrophoblast, trophoblast, and amnion markers in the four TSC types and their parental hPSCs (primed and naïve). Fold changes are normalized to primed hPSCs. Data are presented as mean ± SD (n=3). \**P* < 0.05 compared to primed hPSCs. (D) Proliferation ratios of nTSCs, nbTSCs, ccTSCs, and pdTSCs over a 48-hour interval at passage 13. All TSCs proliferated well, with ccTSC demonstrating slightly slower proliferation rate compared to the other TSCs, though not statistically significant. (E) Schematic representation of protocols for deriving four types of TSCs. nTSC: directly derived from naïve hPSCs; nbTSC: derived from attached naïve hPSC-derived blastoids; ccTSC: generated from primed hPSCs via chemical conversion with HDACi VPA, PD0325901, and LIF, involving an intermediate state of epigenetic reprogramming; pdTSC: induced from primed hPSCs with initial BMP4 and WNTi (IWP2) treatment to prime cells toward the trophectoderm phase. (F) Immunostaining of GATA3 and KRT7 in nTSCs, nbTSCs, ccTSCs, and pdTSCs. Phalloidin and DAPI staining for cytoskeleton and nucleus. Scale bar, 100 μm. (G) Quantitative gene expression analysis of pluripotency, syncytiotrophoblast, trophoblast, and amnion markers in the four TSC types and their parental hPSCs (primed and naïve). Fold changes are normalized to primed hPSCs. Data are presented as mean ± SD (n=3). *P < 0.05 compared to primed hPSCs. (H) Proliferation ratios of nTSCs, nbTSCs, ccTSCs, and pdTSCs over a 48-hour interval at passage 13. All TSCs proliferated well, with ccTSC demonstrating slightly slower proliferation rate compared to the other TSCs, though not statistically significant.

CT differentiation involves interactions among early embryonic cell fates, such as the epiblast and hypoblast within the blastocyst. This process can be recapitulated by forming blastoids from naïve hPSCs, which mimic early axis formation and lineage specification. Prior studies showed that blastoid TE could commit to TSCs^28–30^, though blastoid-derived TSCs (nbTSC) remain incompletely characterized. We generated blastoids from naïve hPSCs cultured in PXGL medium and derived nbTSCs in TSC medium (Figure 1A, S1A). After several days, the attached blastoid outgrowth developed TSC morphology, and a highly homogeneous population emerged upon passage, with marker expression consistent with nTSCs. (Figure 1B, 1C, and S1B).

We generated TSCs from primed hPSCs using two approaches: either through initial BMP4 induction to prime the cells to the TE phase^22^ (primed-state-derived TSCs, pdTSC) or through an intermediate state of epigenetic reprogramming^25^ (chemically converted TSCs, ccTSC) (Figure 1A). The signaling of BMP4 triggers trophoblast induction but also increases mesoderm induction^31,32^. Therefore, for pdTSCs, BMP4 induction was combined with the WNT inhibitor IWP2 to suppress WNT dependent mesoderm induction, guiding cells to a TE-like intermediate phase before stabilizing in a CT-like state in modified TSC medium^33,34^. Conversely, to generate ccTSCs, primed hPSCs were reprogrammed back to an intermediate naïve-like state through epigenetic resetting, using transient histone deacetylase (HDAC) and MEK inhibition with LIF stimulation. HDAC inhibitors, being global epigenetic destabilizers, allow the cells to overcome the epigenetic hurdle that prevents later divergence towards extraembryonic fate. Both pdTSCs and ccTSCs from primed hPSCs showed similar morphology, proliferation rates and expressed trophoblast markers (GATA3, KRT7, GATA2, TFAP2A) as nTSCs and nbTSCs (Figure 1B-D, S1B).

All four TSC types showed downregulation of pluripotent markers and upregulation of key trophoblast markers compared to their parental hPSCs (Figure 1C). Consistent with nTSCs, nbTSCs, and ccTSCs lacked CDX2 expression, while pdTSCs retained low, stable CDX2 levels (Figure S1C). These data indicates that naïve hPSC-derived TSCs, similar to embryo/placenta-derived TSCs, exhibit transcriptome signatures resembling day 8-10 villous CT, distinct from day 5-6 CDX2-expressing TE cells^15^. In contrast, pdTSCs may retain some early TE characteristics. Lastly, We also compared these TSCs with trophoblast-like cells derived from isogenic primed hPSCs treated with BMP4, A83-01, and the FGFR inhibitor PD173074^35,36^ (BAP medium, Figure S1D). BAP cells differed from bona fide TSCs as they lacked robust self-renewal and ceased proliferating after several passages. BAP cells expressed GATA2, TFAP2C, GATA3 and KRT7 (Figure S1B, E) but showed elevated amnion markers expression (Figure 1C), suggesting an amnion-like property distinct from all isogenic TSCs derived.

### Spontaneous Occurrence of Aneuploidy Indicates Active CIN in TSCs

Strikingly, we observed a high incidence of CIN in all these TSCs, a phenomenon not reported in previous studies. Routine karyotyping around passage 10 revealed that over 25% of cells of the 20-40 analyzed cells in each line showed diverse chromosomal aberrations, often involving multiple chromosomes rather than single alterations (Table S1, Figure S2A). The heterogeneity of these aneuploid karyotypes suggests active chromosome mis-segregation in the TSCs, as these non-clonal abnormalities are unlikely to have been inherited from the parental cell lines. Notably, all direct parental cell lines (both naïve and primed hPSCs) were karyotyped immediately before the TSC derivation protocol, and no aneuploidy was detected.

To validate our observation of aneuploidy, we characterized TSCs using fluorescence in situ hybridization (FISH) (Figure 2A), selecting only lines without apparent clonal populations from karyotyping. We chose chromosomes 7, 12, and 18 due to their varying trisomy frequencies during pregnancy: chromosome 7 (frequently detected in placenta), chromosome 12 (rare), and chromosome 18 (viable trisomy) ^37^. Compared to the baseline control—the stable primed hPSC line with an aneuploidy rate of <5% (undetectable by routine karyotyping)—we observed an elevated aneusomy rate in all four TSC types across all three chromosomes starting at passage 4 (Figure S2B), significantly accumulating at high passages (passage>12) (Figures 2B). Using fluorescence-activated cell sorting (FACS) to measure DNA content, we identified aneuploidy by detecting cells with DNA content outside the 2N and 4N ranges and analyzing the S-phase proliferation ratio. Approximately 25% of cells were aneuploid across all four TSC types (Figures 2C, 2D).

**Figure 2.**
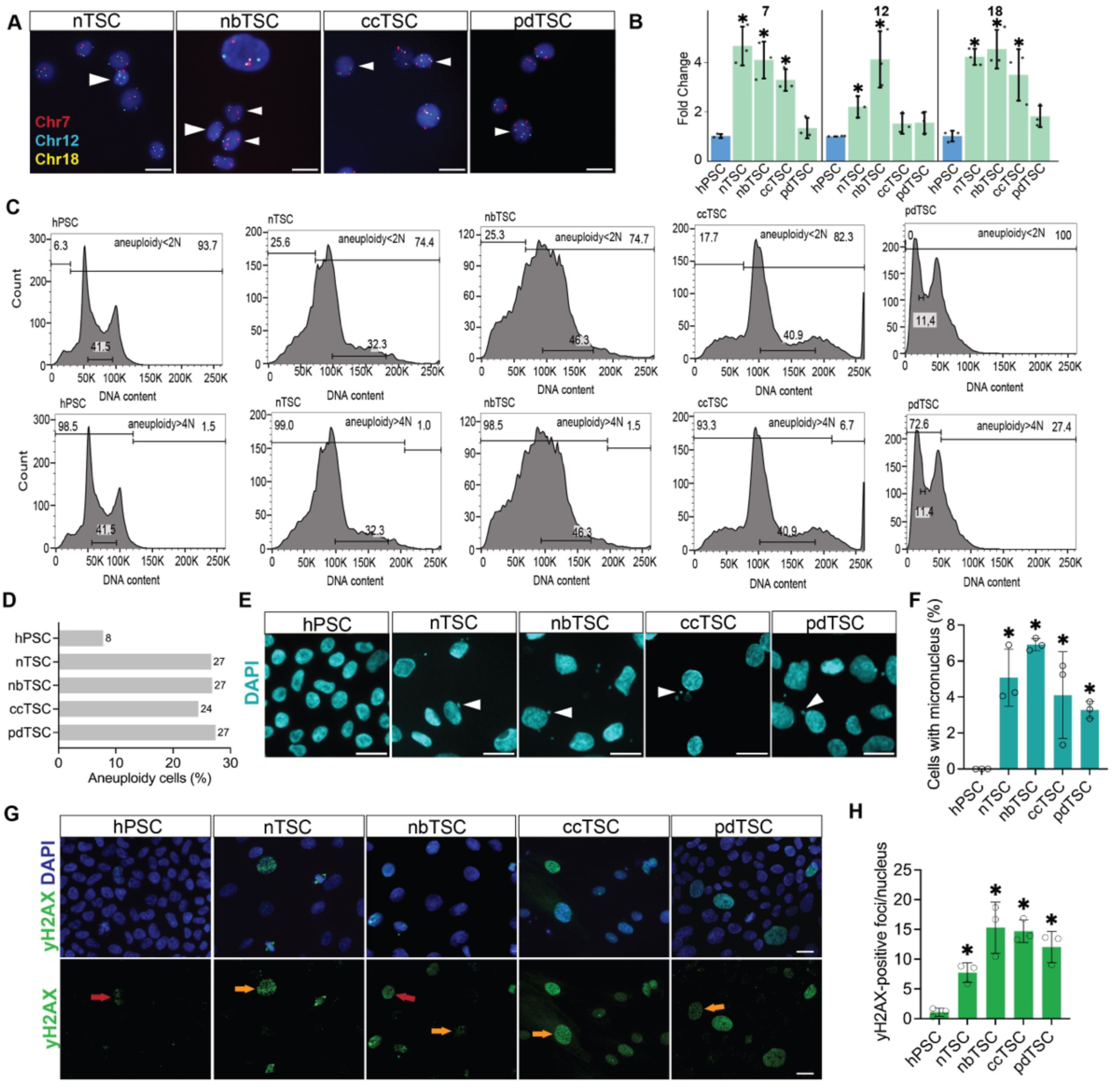
Characterization of chromosomal instability and DNA damage in TSCs. (A) Representative FISH images showing aneuploid cells (white arrows) in nTSCs, nbTSCs, ccTSCs, and pdTSCs, detected using probes for chromosomes 7, 12, and 18. Scale bar, 20 μm. (B) Fold change of aneuploid cell with chromosomes 7, 12, and 18, normalized to the baseline control primed hPSCs. \**P* < 0.05 compared to primed hPSCs. The TSC passage number is from 12 to 16. (C) Flow cytometry analysis of DNA content in TSCs and primed hPSCs. DNA contents < 2N or > 4N indicating aneuploidy. (D) Percentage of aneuploid cells, identified by DNA content outside the 2N and 4N ranges, alongside S-phase proliferation ratio. TSCs exhibit elevated aneuploidy levels compared to parental hPSCs. (E) Representative image of micronuclear DNA (white arrow) in TSCs. DAPI for nuclear staining. Scale bar, 20 μm (F) Percentage of cells with micronuclei in TSCs and primed hPSCs. Mean ± SD (≥500 cells counted per line, n=3). \**P* < 0.05 compared to primed hPSCs. (G) Immunofluorescence staining for γ-H2AX, indicating double-strand DNA breaks, in nTSCs, nbTSCs, ccTSCs, pdTSCs, and hPSCs. Red arrow indicates apoptosis with diffuse expression of γ-H2AX; orange arrow indicates cells with double-strand DNA breaks, dotted expression of γ-H2AX. Scale bar, 20 μm. (H) Quantification of γ-H2AX foci per nucleus in TSCs and primed hPSCs. Mean ± SD of >50 cells per line (n=3). \**P* < 0.05 compared to primed hPSCs.

Since CIN creates extensive karyotype heterogeneity, bulk sequencing and routine karyotyping are insufficient to detect this variability due to limited sampling and technical artifacts. This lack of sensitivity is further supported by our copy number variation (CNV) analysis from bulk DNA whole-genome bisulfite sequencing (WGBS), which failed to detect any aneuploidy in the tested cell lines (Figure S2C). Although WGBS is generally effective for identifying clonal chromosomal gains or losses^38^, it appears ineffective for detecting the non-clonal chromosomal heterogeneity in our TSCs.

Interestingly, in one validated ccTSC line, we observed a clonal effect, with most cells displaying additional copies of chromosome 7 (trisomy 7 or tetrasomy 7, Figure S2D and S2E). These aneuploid cells also showed other chromosomal gains and losses, indicating ongoing CIN (Table S1). The predominance of chromosome 7 additions in this cell line suggests a selective bias or fitness advantage, supported by clinical data showing trisomy 7 as common in human placental samples^39,40^. Additionally, one nbTSC line displayed a substantial chromosome increase (81-95 chromosomes), indicating a clonal tetraploidy status with additional gains or losses (Figure S2F). Derived from a single blastoid, nbTSCs are highly prone to clonal abnormalities. Despite these aberrations, affected cells showed no morphological differences and expressed TSC markers GATA3 and KRT7 (Figure S2E).

Furthermore, similar to CIN cancer cells^41,42^, we observed a high incidence of micronuclei, which form from lagging chromosomes or chromosome fragments that fail to integrate into the primary nucleus after mitosis (Figure 2E and 2F). Furthermore, γH2AX staining for DNA double-strand breaks revealed a significantly increased level of DNA damage in TSCs (Figure 2G and 2H). This damage could be induced either directly through the production of lagging chromosomes or indirectly through replicative stress caused by non-stoichiometric forms of protein complexes involved in replication^43–45^. Together, these findings indicate elevated genomic instability and cellular stress in the TSCs.

Despite this CIN-induced stress, all TSCs tolerated these abnormalities well, continuing to proliferate for over 30 passages in culture. Among the four TSC types, pdTSCs exhibited slightly lower CIN according to FISH and karyotyping.

### Chromosomal Instability and the Presence of Aneuploidy Do Not Compromise the Differentiation Capacity of TSCs

We next conducted functional characterization of TSCs with high CIN to assess their differentiation potential into trophoblast subtypes, including STs and EVTs. All four TSC types showed spontaneous ST differentiation in maintenance culture, with visible cell fusion and some cells expressing ST marker SDC1(Figure S3A and S3B). ST differentiation was also confirmed by detectable human chorionic gonadotropin (hCG) in the culture medium using over-the-counter pregnancy tests (Figure S3C).

Using forskolin, a cAMP agonist, we induced ST differentiation in all TSC types, with syncytia formation observed (Figures 3A and S3D), and nTSC-STs and nbTSC-STs displaying higher SCD1, CGA, CGB, and PSG3 transcription levels, suggesting enhanced ST differentiation efficiency in TSCs derived from naïve origins compared to those derived from primed hPSCs (Figure 3C).

**Figure 3.**
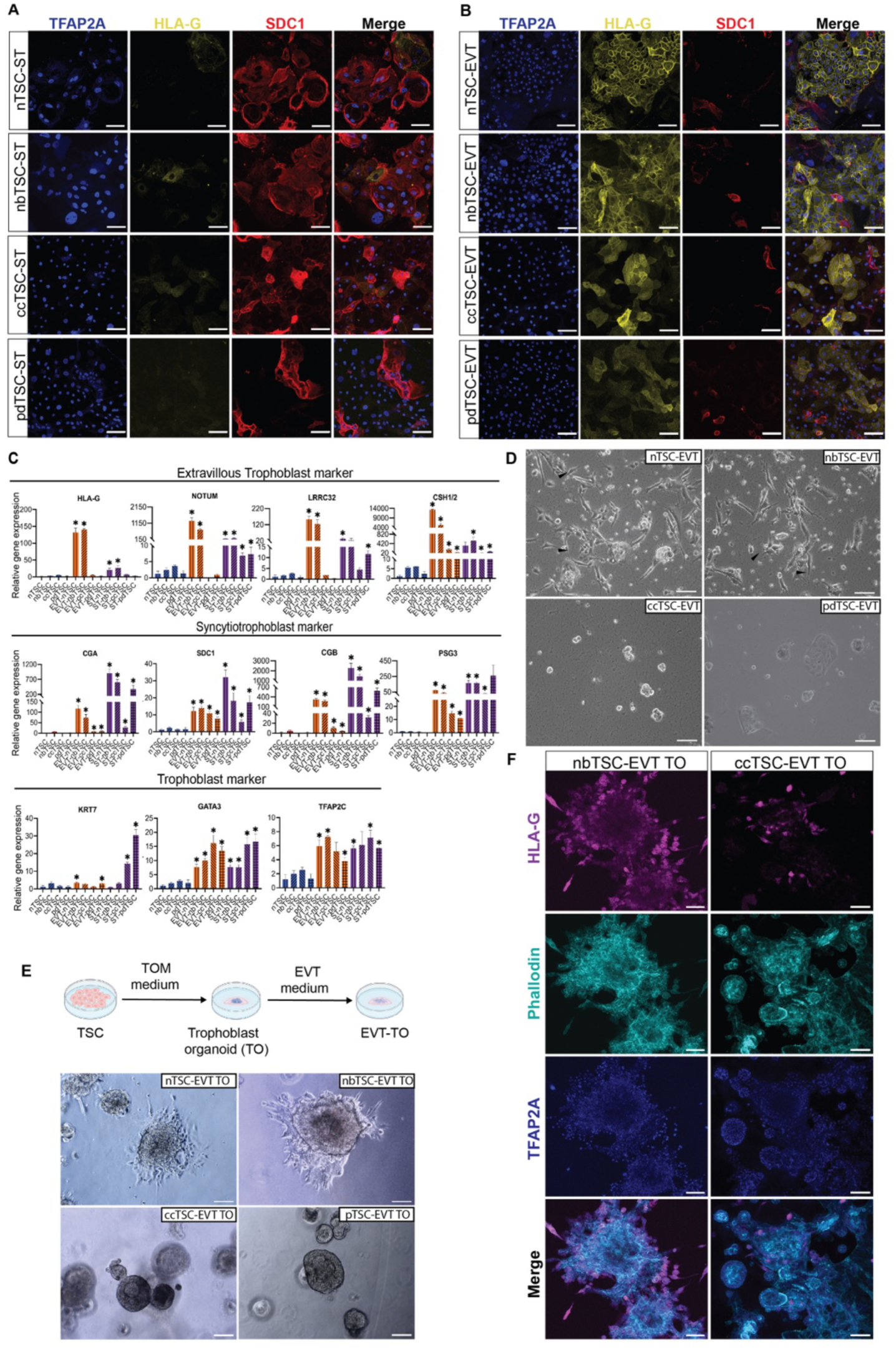
Differentiation capacity of TSCs is uncompromised by CIN. (A) Immunostaining of STs derived from nTSC, nbTSC, ccTSC, and pdTSC. Despite the presence of aneuploidy (passage number >10) and CIN, all four TSC types differentiated into STs expressing the ST marker SDC1 and the trophoblast marker TFAP2A. Scale bar, 100 μm. (B) Immunostaining of EVTs derived from nTSC, nbTSC, ccTSC, and pdTSC. All four TSC types differentiated into EVTs expressing the EVT marker HLA-G, with spontaneous ST differentiation indicated by SDC1 expression. nTSC and nbTSC showed more efficient EVT induction, with a higher nucleus/cytoplasm ratio. Passage number of all TSC types >10. Scale bar, 100 μm. (C) qRT-PCR analysis of ST, EVT, and general trophoblast markers in STs and EVTs derived from all four TSC types. \**P* < 0.05 compared to their parental TSCs. (D) EVTs derived from nTSCs and nbTSCs formed finger-like projections upon replating, while EVTs from ccTSCs and pdTSCs did not display this characteristic. Arrows indicate finger-like projections. Scale bar, 100 μm. (E) EVT differentiation was sufficiently induced only from nTSC-TO and nbTSC-TO, but not from ccTSC-TO and pdTSC-TO. Scale bar, 100 μm. (F) Immunofluorescent staining showed that EVT-TOs derived from nbTSCs fully expressed the EVT marker HLA-G, whereas EVT-TOs derived from ccTSCs did not. Both types expressed the trophoblast marker TFAP2A. Scale bar, 100 μm.

When inducing EVTs, morphological differences emerged between TSCs from naïve and primed origins. After 3 days in NRG1+ culture followed by 3 days without, nbTSCs and nTSCs expressed high protein levels of HLA-G and had a high nucleus-to-cytoplasm ratio (Figure 3B). In contrast, only a subset of pdTSCs and ccTSCs expressed moderate HLA-G levels and displayed a flattened morphology with a lower nucleus-to-cytoplasm ratio (Figure 3B). EVT-specific genes (HLA-G, NOTUM, LRRC32, CSH1/2) were highly upregulated in nbTSC-EVTs and nTSC-EVTs, but barely detectable in pdTSC-EVTs and ccTSC-EVTs (Figure 3C).

Upon splitting on day 6 of induction, nTSC-EVTs and nbTSC-EVTs underwent an epithelial-mesenchymal transition, acquiring a spindle-like morphology, whereas this transition was absent in pdTSCs and ccTSCs (Figure 3D). These observations align with previous findings comparing naïve hPSC-derived TSCs and primed hPSC-derived TSCs through initial BMP4 induction^21^. However, conflicting results from other studies suggest that differentiation capabilities may be cell line-specific, and different signaling pathways may be required for primed TSCs to efficiently generate EVTs^19,25^.

In the 2D differentiation assay, both early-passage TSCs (passage 3, Figures S3E and S3F), where most cells had a normal chromosomal status, and late-passage TSCs (passage 10, Figures 3A and 3B), with higher levels of accumulated aneuploidy, demonstrated similar differentiation capacity. Even TSCs with severe abnormalities, like tetraploid nbTSCs and clonal chromosome 7 additions in ccTSCs, retained full differentiation potential (Figures S2G, S2H). Thus, CIN and high aneuploidy levels seem well tolerated in these trophoblast lines, with no compromise in cellular function, consistent with previous findings of significant aneuploidy in invasive EVT cells^7^.

### Trophoblast Stem Cells Derived from Primed hPSCs Exhibit Moderate EVT Differentiation Capacity

Previous studies have shown that naïve hPSC-derived TSCs can generate trophoblast organoids (TOs) similar to those from primary TSCs in tissue architecture, hormone secretion, and self-renewal^46,47^. However, the ability of TSCs derived from primed hPSCs to form TOs remains unexplored. We seeded four TSC types in 3D Matrigel droplets with trophoblast organoid medium, as described by Turco et al^48,49^. Rapid cell expansion occurred in all TSCs, forming complex organoids within a week, though primed hPSC-derived TOs showed greater size variation (Figure S4A). Organoid growth rates were higher in nTSC and nbTSC, producing larger organoids than those from pdTSC and ccTSC (Figure S4B). Within two weeks, the organoids reached 200–300 μm and were passaged by mechanical and enzymatic dissociation.

To further assess their differentiation potential, TOs larger than 100 μm were seeded into Matrigel and treated with EVT induction medium. nbTSC-TOs and nTSC-TOs attached and produced outgrowing EVT cells with spindly, elongated morphology, staining positive for HLA-G (Figure 3E and 3F). In contrast, most ccTSC-TOs and pdTSC-TOs expanded without attaching in EVT medium, with only a small fraction of attached organoids displaying HLA-G-positive cells (Figure 3E and 3F). Less than 1% of these TOs attached, typically at the Matrigel droplet edge, with most cells lacking EVT morphology.

Given the moderate EVT differentiation capacity of ccTSCs and pdTSCs, we conclude that TSCs derived from primed hPSCs have significantly lower differentiation potential compared to naïve hPSC-derived TSCs. This supports previous claims that primed hPSCs are less capable of generating bona fide trophoblast cells, while naïve hPSCs offer a more robust model for studying placental dynamics due to their efficient induction of downstream trophoblast subtypes.

### Trophoblast Stem Cells Derived from Naïve Origins Exhibit Closer Transcriptome and Methylome Profiles to Cytotrophoblasts

To further assess global gene expression and DNA methylation landscapes in these TSCs, we performed RNA-seq and WGBS on cells at passages 9–12. All four isogenic TSC types clustered closely in transcription profiles, distinct from BAP cells, and from both primed and naïve hPSCs (Figure 4A and 4B). This separation was further supported by differential expression of pluripotency and amnion markers (Figure 4C). Among the TSCs, nbTSC and nTSC grouped together, while pdTSC and ccTSC clustered more closely (Figure 4A). Comparisons to first-trimester CTs, blastocyst-derived TSCs (TSblast), and placenta tissue-derived TSCs (TSCT)^10^ revealed that nTSCs and nbTSCs were more similar to TSblast and TSCT, with blastoid-derived nbTSC clustering closest to TSblast (Figure 4B).Additionally, pdTSC and ccTSC grouped with other primed-derived TSCs from different parental hPSCs, while nbTSC and nTSC clustered with other naïve-derived TSCs (Figure S5A), indicating that our isogenic TSCs share profiles consistent with previous studies^13,19,22,23,25^.

**Figure 4.**
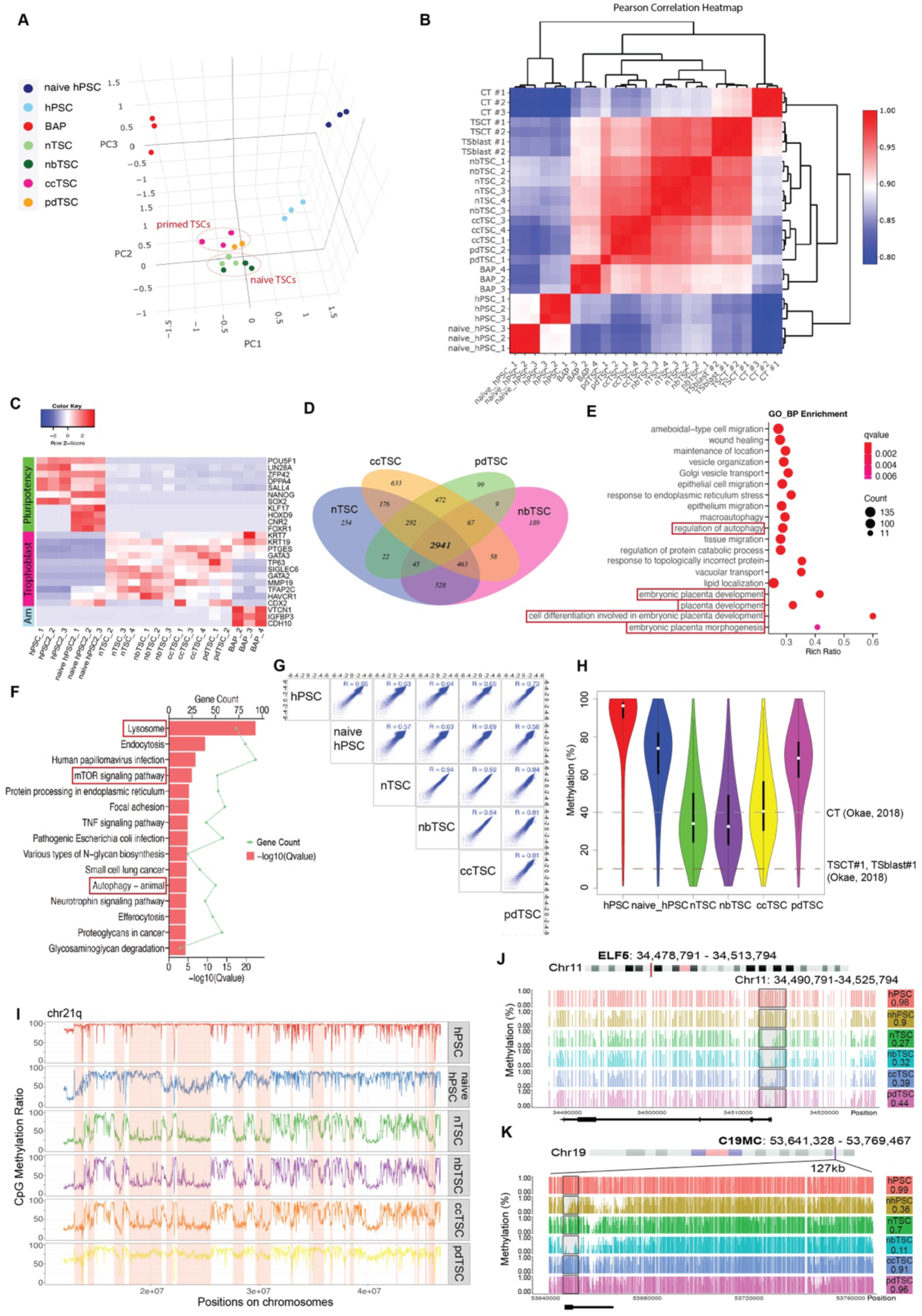
Comparison of four types of TSCs through transcriptome and DNA methylome analyses. (A) PCA showing that naïve- and primed-derived TSCs closely cluster together but are distinct from hPSCs, naïve hPSC, and BAP cells. (B) Heatmap of Pearson correlation between TSCs, hPSCs, CT from primary placenta, and CT- and blastocyst-derived TSCs (TSCT and TSblast)^10^. The dendrogram hierarchy shows that naïve-derived TSCs (nTSCs and nbTSCs) are more closely correlated with primary TSCs. (C) Heatmap of Z score-transformed read counts for specific markers of pluripotency, trophoblast, and amnion cell types in TSCs, primed and naïve hPSCs and BAP cells. (D) Differential gene expression analysis identified 2,941 commonly upregulated genes in the four TSC types during differentiation from parental primed hPSCs. (E) Gene Ontology (GO) biological process analysis of the 2,941 upregulated genes showing significant enrichment in autophagy regulation, placental development, and cell differentiation involved in embryonic placental development. (F) KEGG pathway analysis of the 2,941 upregulated genes showing significant enrichment in lysosome function, mTOR signaling, and autophagy pathways. (G) Spearman correlation analysis comparing DNA methylation levels across TSCs and hPSCs (naïve and primed). WGBS data were analyzed in 10-kb windows. (H) Violin plots showing DNA methylation levels in placenta-specific PMDs. Horizontal dashed lines indicate methylation levels of CT, TSCT, and TSblast^10^. White dots represent the median, and boxplot lines are overlaid within the violin plots. (I) DNA methylation patterns along the long arm of chromosome 21 in TSCs, primed and naïve hPSCs. The vertical axis shows CpG methylation ratio, with PMDs highlighted in pink. (J) DNA methylation patterns at the ELF5 locus. The ELF5 promoter region is within rectangle, with corresponding methylation levels indicated on the right. (K) DNA methylation patterns at the C19MC locus. The C19MC DMR (GRCh38: 19 53647735-53654142) is within rectangle, with corresponding methylation levels indicated on the right.

During the derivation process from parental primed hPSCs, 2,941 genes were upregulated in TSCs, enriched in autophagy, placental development, and embryonic placenta differentiation (Figures 4D–4F). Interestingly, genes downregulated in TSCs relative to primed hPSCs were consistently linked to genomic stability and maintenance, including processes such as DNA repair, replication, chromosomal organization, remodeling, and double-strand break repair (Figure S5B).Pairwise comparisons of the four TSC types (Figures S5C–S5H) showed that naïve hPSC-derived TSCs consistently expressed higher levels of certain zinc finger proteins, including PEG3, a paternally expressed imprinted gene associated with placental growth (Figures S5E–S5H). Conversely, IGF2, another paternally imprinted gene linked to placental growth, was most elevated in primed-derived TSCs (Figures S5E–S5H). The discrepancy between the expression of these two critical imprinted genes in TSCs from different PSC origins is worth further investigation.

All TSCs showed unique DNA methylation features, including global hypomethylation across whole-genome and promoter CpGs (Figure S6A, S6D). nTSC, nbTSC, and ccTSC had nearly identical methylation patterns (R > 0.92), while pdTSC exhibited a slightly lower correlation (R = 0.81–0.84, Figure 4G), reflecting the distinct reprogramming steps in naïve hPSCs and intermediate phase of epigenetic reprogramming in ccTSC derivation. Meanwhile, pdTSC showed a stronger positive linear correlation with primed PSCs, indicating a greater retention of epigenetic markers from the parental cells. This less efficient epigenetic resetting and comparatively lower levels of genomic hypomethylation in pdTSCs may contribute to slightly lower CIN in pdTSCs. All TSCs exhibited placenta-specific partially methylated domains (PMDs), previously shown to cover 37% of the genome and remain stable throughout gestation and across individuals^50^ (Figure 4H). nbTSC, nTSC, and ccTSC had PMD levels comparable to first-trimester CTs, while pdTSC showed higher PMD levels (Figure 4H). Interestingly, we did not observe the pronounced hypomethylation typically seen in primary tissue- and blastocyst-derived TSCs (TSCT#1 and TSblast #1). However, PMD patterns were consistently maintained across all four TSC types, with nbTSC, nTSC, and ccTSC showing more similar patterns (Figure 4I, S6B and S6C).

Moreover, the ELF5 promoter, which is characteristically hypomethylated in trophoblasts^51^, was hypomethylated in TSC lines, confirming successful programming to trophoblast fate (Figure 4J). In line with Kobayashi et al.’s findings that the primate-specific miRNA cluster C19MC, a placenta-specific paternally expressed imprinted gene, is active in naïve TSCs but epigenetically silenced in BMP-pretreated primed TSCs, our data reveal that the C19MC DMR, which serves as the promoter for this miRNA cluster, is more demethylated in naïve-derived TSCs, with nbTSC showing the greatest hypomethylation. In contrast, both primed-derived TSCs remained highly methylated (Figure 4K). The correlation between promoter methylation and gene expression revealed that most genes lost methylation and were upregulated during TSC derivation from hPSCs, such as PEG3 in the naïve hPSC-derived TSCs (Figure S6E). In addition, PEG3 and IGF2 displayed distinct methylation patterns in their DMRs, corresponding to the observed gene expression difference for PEG3 but not for IGF2 (Figure S6F and S6G)

In summary, our transcriptome and methylome data from isogenic TSCs derived using different methodologies confirm that naïve cells give rise to more bona fide trophoblast fates. While primed-derived TSCs are sufficiently similar to cytotrophoblasts, they retain intrinsic differences that may contribute to their compromised functionality.

### Less Stringent Cell Cycle Safeguard and Elevated Autophagy Contribute to CIN in TSCs

Unlike differentiated somatic cells, hPSCs frequently acquire CIN^52,53^. However, the proportion of aneuploid cells in hPSC cultures remains low, likely due to compromised cellular fitness or cell competition^54^. In contrast, TSCs tolerate these aberrations to a greater extent, allowing them to proliferate and differentiate without obvious differences from their normal counterparts. This discrepancy in CIN tolerance may be attributed to intrinsic characteristics of TSCs. Thus, we next sought to understand the molecular basis for the formation and tolerance of aneuploidy and CIN in TSCs.

Compared to the parental primed hPSCs, we found that key genes involved in cell cycle process were downregulated in TSCs, particularly those associated with cohesion loading, kinetochore assembly, and the spindle assembly checkpoint (SAC, Figure 5A). In cancer, reduced expression of cohesion-related genes has been linked to CIN^55^. Knockdown of cohesion complex components such as RAD21 and essential SAC genes like MAD2 has been shown to induce CIN in both cancer cells and mice^56–60^. Among the most significantly downregulated genes of this category in TSCs compared to their parental cells were MAD2 (MAD2L1 and MAD2L2) and the complex component BUB1B, suggesting SAC inactivation a potential cause for active chromosome mis-segregation in TSCs (Figure 5B). Additionally, RAD21 was notably downregulated in TSCs but not in pdTSCs, which pinpoints another potential cause for CIN and may explain the lower aneuploidy observed in the latter (Figure 5B).

**Figure 5.**
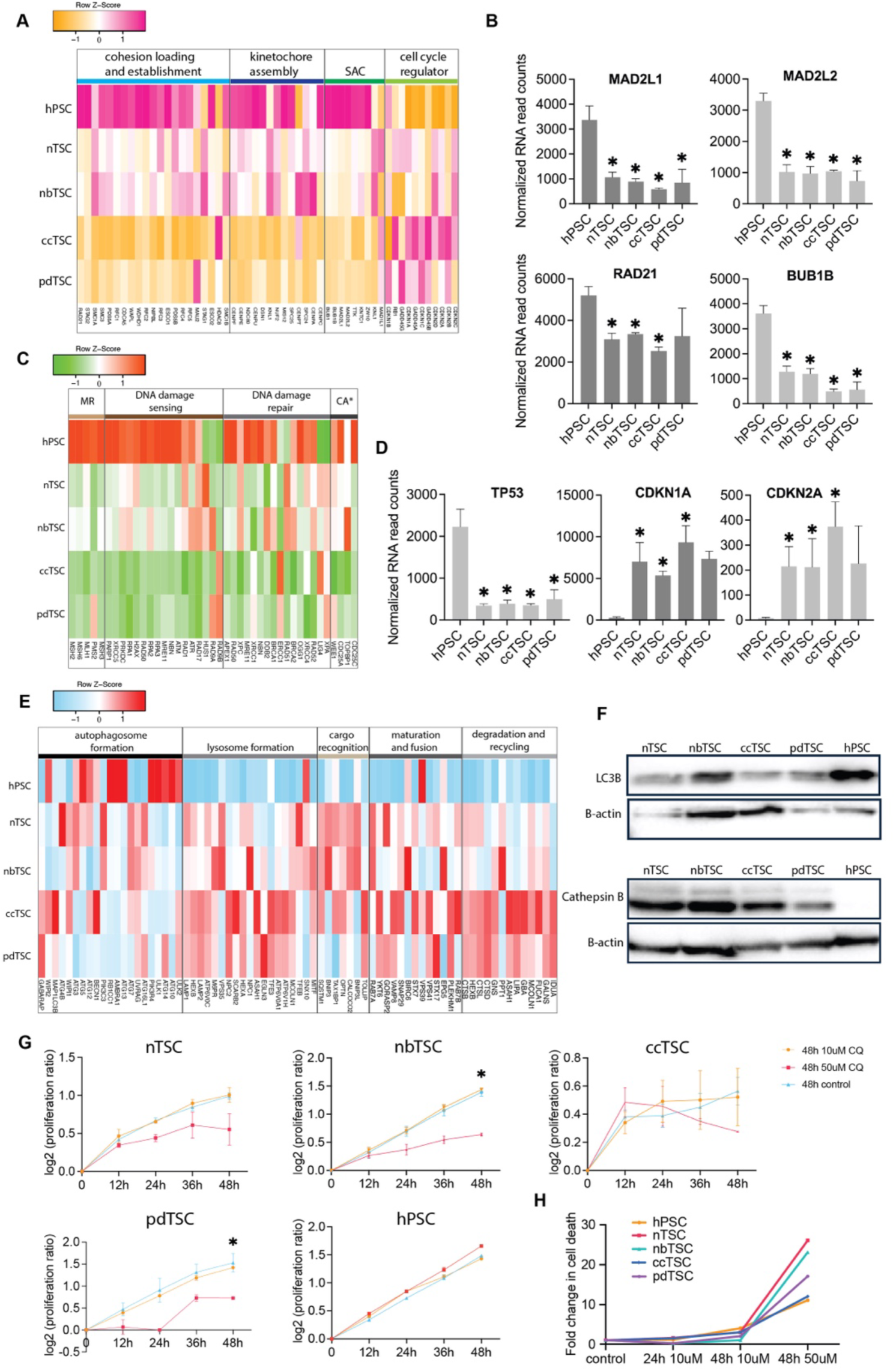
Reduced cell cycle safeguards and elevated autophagy in chromosomally unstable TSCs. (A) Heatmap of Z score-transformed read values for genes involved in key processes regulating the cell cycle. Compared to primed hPSCs, TSCs show overall downregulation in cohesion loading and establishment, kinetochore assembly, and spindle assembly checkpoint (SAC) genes, with upregulation in cell cycle regulators specific to G1 inhibition. (B) Normalized RNA read counts of SAC genes (MAD2L1, MAD2L2, and BUB1B) and the core cohesion complex gene, RAD21. \**P* < 0.05 compared to primed hPSCs. (C) Heatmap of Z score-transformed read values for genes involved in DNA and genomic damage repair. Compared to parental hPSCs, TSCs display overall downregulation in mismatch recognition (MR), DNA damage sensing, DNA repair, and checkpoint activation (CA) pathways. (D) Normalized RNA read counts of the TP53, CDKN1A, CDKN2A genes. \**P* < 0.05 compared to primed hPSCs. (E) Heatmap of Z score-transformed read values for genes involved in autophagy-related processes. TSCs show no apparent change in autophagosome formation compared to parental hPSCs; however, there is overall upregulation in lysosome formation, cargo recognition, autophagosome maturation and fusion with lysosomes, as well as degradation and recycling processes. (F) Western blot analysis of LC3B and Cathepsin B expression in TSCs and primed hPSCs. B-actin as a loading control. (G) Changes of proliferation ratio of TSCs and primed hPSCs following chloroquine (CQ) treatment at a 48h interval. Data presented as mean ± sd (n=3). \**P* < 0.05, 48h 50uM CQ compared to 48h control. (H) Fold change in cell death measured by flow cytometry using Sytox staining after CQ treatment in TSCs and primed hPSCs.

Despite elevated DNA damage and CIN, TSCs did not exhibit a corresponding increase in DNA repair activity; in fact, transcript levels of genes involved in mismatch repair, DNA damage response, and repair pathways were downregulated (Figure 5C). Among them, the tumor suppressor p53 was significantly reduced in all TSC lines compared to hPSCs, regardless of the genotoxic stress from CIN (Figure 5D). This suggests that low p53 levels play a central role in the permissive environment for chromosomal abnormalities in TSCs. While p53 levels increase in the placenta during complicated pregnancies, facilitating trophoblast apoptosis under stress^61^, maintaining relatively low p53 levels in TSCs may be necessary for normal trophoblast function, even if it compromises genomic integrity. Interestingly, unlike tumorigenesis, where cell cycle regulators are typically downregulated to allow unchecked proliferation^62–65^, TSCs maintain better cell cycle control despite high genomic instability. This is likely achieved by upregulating cell cycle regulators, specifically CDK and G1 inhibitors (Figure 5A). Among these regulators, the key senescence genes P16 (CDKN1A) and P21(CDKN2A) are the most significantly upregulated, potentially preventing them from transforming into malignant cells (Figure 5D).

Interestingly, we observed a massive upregulation of most autophagy-related genes in TSCs, especially those involved in lysosome formation, fusion, and degradation, though autophagosome initiation genes were not notably upregulated (Figure 5E). This pattern was confirmed at the protein level: the autophagosome initiation marker LC3B-II was not elevated in TSCs, whereas the lysosomal degradation marker, the cysteine protease cathepsin B (CTSB), was highly expressed in TSCs but undetectable in the parental primed hPGCs (Figure 5F). This suggests a metabolic shift essential for the survival of chromosomally unstable cells, as aneuploidy-induced protein imbalances cause proteotoxic stress, triggering autophagy to degrade and recycle damaged proteins. In a Drosophila model, autophagy was shown to be activated in CIN cells and necessary for their survival^66^. In cancer, autophagy plays a dual role, preventing cancer expansion by maintaining cellular homeostasis but also supporting cell survival under stress^67^. Previous studies have shown that inhibiting autophagy facilitates the development of aneuploid mouse epiblasts, suggesting that autophagy helps eliminate aneuploid cells through apoptosis^68^. However, autophagy likely has more complex and context-dependent roles in early embryogenesis. The significant upregulation of autophagy pathways in TSCs indicates that autophagy serves as a survival mechanism, enabling these cells to tolerate high levels of genomic instability by degrading toxic components. This was further confirmed by inhibiting autophagy with chloroquine (CQ) at autophagosome-lysosome fusion stage (Figure S7A). CIN TSCs were more affected by this inhibition compared to the relatively stable primed hPSCs. Treatment with 50 µM CQ effectively reduced cell proliferation and increased cell death in TSCs, while hPSCs remained largely unaffected (Figure 5G, S7B). Lower CQ doses were less effective, likely due to the already high basal levels of autophagy in TSCs. Our findings align with Tang et al., who reported increased basal autophagy in aneuploid MEFs following chromosomal mis-segregation and noted that aneuploid cells were more sensitive to autophagy inhibition^69^.

### Identification of CIN in Developing Human Placentas

The next obvious question is whether the CIN and tolerance of aneuploidy observed in TSCs truly reflect what occurs in vivo during placental development. The widely accepted view that aneuploidy is rare in the placenta and negatively impacts pregnancy outcomes warrants reconsideration, as previous methods for analyzing placental samples, such as FISH, karyotyping, and bulk CNV analysis by qPCR, CGH, or sequencing, have limited resolution. FISH and karyotyping examine only a small number of cells, making them prone to artifacts, especially when heterogeneous aneuploidy is present in a minority. Additionally, these methods often require cell culture to obtain metaphase spreads, where suboptimal in vitro conditions may introduce stress that favors the survival of normal cells, potentially eliminating aneuploid cells outside the placental niche. CIN results in substantial karyotypic heterogeneity, rendering bulk CNV methods inadequate for detecting this variation.

We hypothesized that high CIN levels and prevalent aneuploidy also occur in placental trophoblast cells. To test this, we analyzed single-cell RNA sequencing (scRNA-seq) and single cell multiome data from placentas derived from healthy pregnancies^70,71^. Using a similar approach to our analysis of human embryo data^72,73^, we examined plate-based Smart-seq data from eight first- and second-trimester placentas. To reduce false negatives due to transcriptional heterogeneity, we based our analysis on UMAP clusters (Figure S8A and S8B). We excluded marker-defined STs from the analysis due to their multinucleation, which disrupts RNA-based CNV detection accuracy. Aneuploidy was most prevalent in certain CT clusters, with moderate levels in EVTs (Figure S8B). However, the sequencing quality was suboptimal, and many clusters did not meet quality control criteria based on gene count per cell, reducing data reliability (Table S2). The only cluster that passed quality control was 24-week EVT cluster, where 9.2% aneuploidy was detected (Figure S8C).

We analyzed scMultiome data from 12 healthy placentas, with six from early (6–9 weeks) and six from term pregnancies (38–39 weeks)^71^, where CTs and STs were included. By using single nuclei, the data avoided inaccuracies associated with syncytia in STs. While scRNA-seq is limited by factors like gene compensation and karyotypic heterogeneity, scATAC-seq better predicts CNV by reflecting directly the DNA content.^74–76^. Using CNV predictions from EpiAneuFinder^76^, we mapped read count data from single-nuclei ATAC-seq to detect genome-wide copy number alterations (CNAs), allowing us to assess CNA heterogeneity at the single-cell level (Figure 6A and 6B).

**Figure 6.**
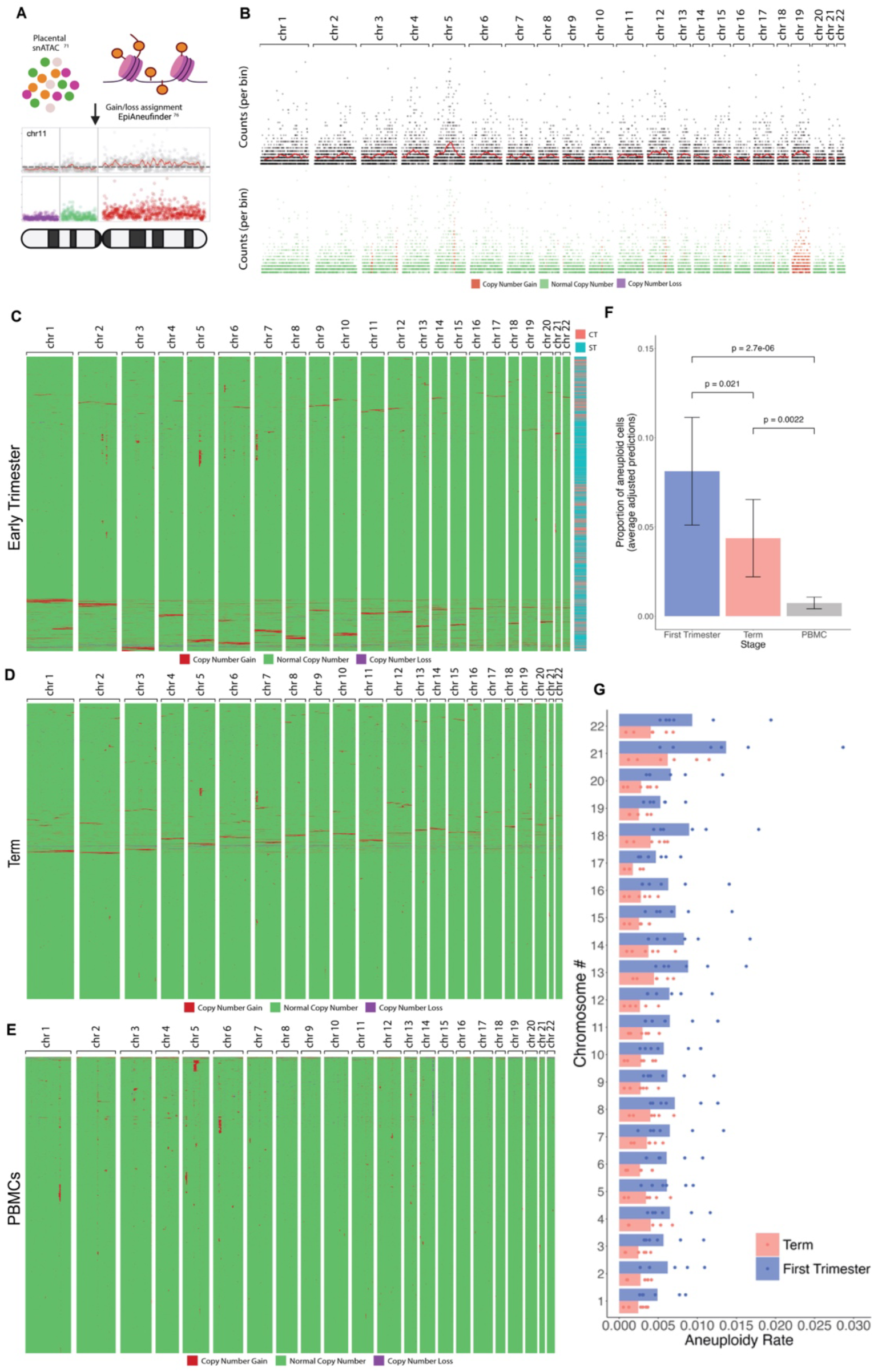
Detection of copy number alterations and aneuploidy in human placenta. (A) Schematic representation of the strategy for using EpiAneuFinder on available human placenta snATAC-seq data to identify copy number alterations (CNAs) at single-cell resolution. Segmented chromosomes are assigned a state of loss, normal, or gain based on the read count fold change relative to the genome-wide mean. (B) Representative count plotting of a single cell displaying a whole chromosome gain for chromosome 19. (C) CNA karyogram for CT and ST nuclei derived from a representative first trimester placenta. Each row on the y-axis represents a cell, and the x-axis shows chromosomal locations. Normal copy numbers are displayed in green, gains in red, and losses in purple. (D) CNA karyogram for CT and ST nuclei from a representative term placenta. (E) CNA karyogram for PBMCs from a healthy adult individual. (F) Average adjusted predictions of the proportion of aneuploid cells in each sample type, based on a binomial generalized linear mixed model (GLMM) with tissue donor as a grouping factor. Error bars indicate 95% confidence intervals, and p-values reflect pairwise comparisons between average adjusted predictions. (G) Percentage of aneusomic cells per chromosome in first trimester (n=6) and term placentas (n=6). The rate of aneuploidy represents the frequency at which each individual chromosome had at least 60% of bins classified as a CNV by EpiAneuFinder across all cells for each sample.

Strikingly, compared to human peripheral blood mononuclear cells (PBMCs) with similar sequencing depth, which showed minimal CNA, placental samples exhibited significantly higher CNA levels, primarily involving whole chromosome gains or losses (Figure 6C-F and S9B-D). Whole chromosome gains were notably more frequent than losses, likely because the method is less sensitive to detecting losses, resulting in possible false negatives^76^. Alternatively, cells with whole chromosome losses may be less viable, limiting their presence in the placenta. No significant CNA differences were observed between CTs and STs, as cells with CNAs were distributed evenly across both types (Figure 6C).

It is noteworthy that the fragment/cell count in this dataset was approximately five-fold lower than the recommended optimal level^76^ (Figure S9A). When the optimal fragment count was downsampled five-fold, EpiAneuFinder maintained high precision in CNV detection (precision scores >0.8), but the recall rate dropped to ∼0.4, increasing the chance of false negatives with lower sequencing depth^76^. Therefore, the actual rate of CNA in the placenta is likely higher than detected. Even with this limitation, and by counting only aneuploid cells with more than 60% CNA in a single chromosome, we found an average of 8% aneuploid trophoblast cells in first-trimester placentas, compared to 0.7% in PBMCs (Figure 6C and 6E). Interestingly, this percentage dropped significantly in term placentas (4.4%, Figure 6D and 6F), suggesting that the placental environment may become more selective against aneuploid cells as pregnancy progresses.

We also observed extensive heterogeneity in aneuploidy, with chromosomal changes detected across all 22 chromosomes, indicating active CIN rather than clonal aneuploidy (Figure 6C-E). Interestingly, the most frequent changes involved chromosomes associated with viable trisomies (13, 18, 21), suggesting that these trisomies may confer a higher degree of cellular fitness (Figure 6G).

## Discussion

In this study, we present compelling evidence that CIN is an inherent feature of TSCs and trophoblast lineages in human placentas. By deriving isogenic TSC lines from primed and naïve human PSCs and blastoids, we minimized cell line and methodology-specific biases toward CIN. Our study also directly compares methodologies for generating TSC lines, confirming that naïve PSCs are preferable for producing bona fide trophoblast lineages. Strikingly, we observed prevalent aneuploidy across all TSC types, regardless of derivation method. The extensive heterogeneity in karyotypes suggests active mis-segregation in these cells, likely due to the downregulation of key cell cycle safeguard genes. This hypothesis is further supported by analysis of healthy placentas, where we detected significant populations of heterogeneous aneuploid cells. Despite high levels of aneuploidy, chromosomal alterations, and DNA damage, TSCs retained their proliferative and differentiation capacities, showing remarkable resilience to genomic instability. Our data also reveal an intrinsic role of autophagy in tolerating aneuploidy and genomic alterations, suggesting it as a protective mechanism in these cells.

Our previous work, along with others, has demonstrated that human embryogenesis is particularly prone to CIN and aneuploidy, especially in the early stages, but these abnormalities are gradually eliminated during differentiation ^73,77–79^. This is a major cause of the low efficiency of human reproduction, with nearly half of fertilized eggs failing before the first trimester. We showed that TE fate is more tolerant of aneuploidy, raising the question of whether this dynamic persists throughout placental development^73^. With the recent development of more bona fide trophoblast models, we now have an excellent platform to investigate these questions.

Our findings challenge the traditional view that placental aneuploidy is solely pathological and linked to adverse pregnancy outcomes. Instead, our data suggest that CIN and aneuploidy are inherent features of trophoblasts, potentially serving physiological roles in normal placental development. These abnormalities may play a physiological role in normal placental development, as they are part of a package that includes accelerated growth rates, which may come at the cost of compromised cell cycle safeguards. Supporting our hypothesis, Weier et al. have also detected high levels of abnormalities in the human placental cells cultured in vitro^6,7^. Two recent bulk sequencing studies highlighted significant genomic mutations, projecting the placenta as a “dumping ground” for developmental anomalies^77,78^. Our analysis of healthy placental tissue, using single-cell resolution, provides direct evidence of widespread CIN.

While CIN is generally detrimental to cellular fitness in normal tissues, it paradoxically serves as a driver of tumor evolution, with 90% of solid tumors exhibiting aneuploidy and CIN as hallmarks^79,80^. Based on our findings, we speculate that CIN may similarly provide trophoblasts with a survival advantage by enabling genomic flexibility, an adaptation that could be an evolutionary mechanism specific to humans, given our highly invasive placentas. Understanding how the placenta differs from uncontrolled cancer cells could shed light on ways to control abnormal cell behavior. The upregulation of cell cycle regulators found in our study may contribute to the placenta’s ability to maintain control and prevent cells from becoming cancerous. Similar to cancer cells, autophagy plays a critical role in tolerating abnormalities by managing the proteotoxic stress caused by genomic imbalance during fetal and placental development. This is further supported by studies showing that impaired autophagy contributes to pregnancy failure^81,82^. The insights revealed here could pave the way for therapeutic interventions to address placenta-related pregnancy complications. We hope that this comprehensive reevaluation of placental CIN will lead to new directions in the study of the placenta, which is a highly unique and essential cancer-like organ.

## Limitations of the study

While we thoroughly characterized our PSC-derived TSCs and confirmed their similarities to primary trophoblast cells in transcriptional, epigenetic, and differentiation capacities, a key limitation of this study is its reliance on in vitro trophoblast models, which may not fully capture the complexity of CIN or aneuploidy in vivo. Future studies should aim to validate these findings in vivo. Since aneuploidy in embryogenesis is primate-specific, non-human primates (NHPs) would be ideal models, as they more closely resemble human placental development than rodents, which have distinct placentation processes.

The broader role of CIN in the placenta remains unclear. While our data suggest that CIN is a normal feature of trophoblasts, particularly in early pregnancy, its contribution to placental function in later gestation or in pregnancy complications is unknown. Future studies should investigate the functional consequences of CIN in different trophoblast subtypes and its role in placental diseases, as well as determine the extent to which CIN is tolerated in normal pregnancies.

## Resource availability

Further information and requests for resources and reagents should be directed to and will be fulfilled by the lead contact, Min Yang.

## Material and availability

All stable reagents generated in this study are available upon request from the lead contact.

## Data and code availability

All sequencing data generated in this study have been deposited in the NCBI BioProject database and are publicly available as of the date of publication. The accession numbers are GSE280832 (RNA-seq fastq.gz file) and PRJNA1181850 (WGBS-seq fastq.gz file). This paper also utilizes publicly available datasets, including GSE247038 (placenta scMultiome) and GSE89497 (placenta single-cell Smart-seq2). PBMC scMultiome datasets were acquired from 10x Genomics sources webpage.

Any additional information required to reanalyze the data reported in this paper is available from the lead contact upon request.

## Methods

### Conventional hPSC culture

Primed hPSCs (registered RUES2 human embryonic stem cell line, NIHhESC-09-0013) were obtained from Dr. Ali Brivanlou at Rockefeller University. Cells were cultured in mouse embryonic fibroblast-conditioned HUESM medium (MEF-CM)^83^, supplemented with 20 ng/mL basic fibroblast growth factor (bFGF; R&D, 233-FB-500) or mTeSR™ medium (STEMCELL Technologies, 85850). Culture medium was refreshed daily. Tissue culture dishes were coated with Geltrex™ (Gibco, A1413202), applied overnight at 4°C, and subsequently incubated at 37°C for at least 20 minutes before cell plating. Cells were passaged using Accutase™ (STEMCELL Technologies, 7922) and replated in MEF-CM or mTeSR™ containing 10 μM ROCK inhibitor Y-27632 (STEMCELL Technologies, 72308). G-banding karyotyping was performed at Cell Line Genetics to verify chromosomal status prior to initiating experiments.

### Naïve hPSC reprogramming and culture

Reprogramming of the hPSCs to a naïve state was performed according to a previously established protocol with minor modifications^27^. In brief, primed RUES2 were dissociated with Accutase and plated on a gelatin-coated plate with an MEF layer at a density of 10,000 cells/cm. Cells were cultured in 5% CO₂ and 5% O₂ at 37°C. The same medium was refreshed the following day, allowing cells to form colonies. Two days after plating, the medium was changed to N2B27 medium consisting of 50% DMEM-F12 (Gibco, 11330057), 50% Neurobasal (Gibco, 21103049), 2 mM L-glutamine (Gibco, A2916801), 55 μM β-mercaptoethanol (Gibco, 21985023), 0.5× N2 supplement (Gibco, 17502048), and 0.5× B27 supplement (Gibco, 17504044), supplemented with 10 ng/mL human leukemia inhibitory factor (LIF; STEMCELL Technologies, 78055), 1 μM PD0325901 (MEKi; STEMCELL Technologies, 72184), and 1 μM valproic acid (VPA; HDACi; STEMCELL Technologies, 72292). After three days in this medium, it was switched to PXGL medium (N2B27 medium supplemented with 1 μM PD0325901, 2 μM XAV939 [STEMCELL Technologies, 72674], 2 μM Gö6983 [STEMCELL Technologies, 72462], and 10 ng/mL LIF), refreshed daily for an additional 8-10 days before passaging. Naïve hPSCs were passaged on MEF plates using Accutase™ dissociation every 3–5 days in PXGL medium, supplemented with 10 μM Y-27632 and 1% Geltrex™ at a ratio of 1:4 to 1:6. Cells from passages 6–15 were used for downstream experiments. G-banding karyotyping was performed at Cell Line Genetics to verify chromosomal status before initiating experiments.

### Blastoid differentiation

Naïve hPSCs were differentiated to form blastoids according to a previously established protocol with minor modifications^30^. In brief, naïve PSCs were dissociated into single cells using Accutase, and MEFs were excluded by sequential attachment on gelatin-coated plates for 45 minutes. Unattached human naïve hPSCs were seeded on Aggrewells™ 400 (STEMCELL Technologies) at a density of ∼80 cells/microwell in N2B27 medium supplemented with 10 μM Y-27632. Cells were cultured in 5% CO₂ and 5% O₂ at 37°C. On the following day, the medium was changed to N2B27 medium supplemented with PALLY, consisting of 1 μM PD0325901, 1 μM A83-01 (STEMCELL Technologies, 72024), 0.5 μM 1-Oleoyl Lysophosphatidic Acid (LPA; STEMCELL Technologies, 72694), 10 ng/mL LIF, and 10 μM Y-27632. Cell aggregates were cultured in this medium for 48 hours with daily medium change. The cell aggregates were then cultured in N2B27 medium supplemented with LY (0.5 μM LPA and 10 μM Y-27632) for another 48 hours with daily medium change.

### Culture of BAP cells derived from primed hPSCs

Primed hPSCs were dissociated into single cells using Accutase and plated on Geltrex-coated tissue culture plates at a density of 12,000 cells/cm² in MEF-CM supplemented with bFGF. On the following day, the medium was replaced with MEF-CM supplemented with 10 ng/mL BMP4, 1 μM A83-01, and 0.1 μM PD173074. The medium was refreshed daily for up to eight days.

### Derivation of TSCs from primed hPSCs through TE-like cells (pdTSC)

Derivation of pdTSC was preformed according to a previous two-step protocol with modifications^22^. In brief, hPSCs were dissociated using Accutase and plated onto collagen IV (5 μg/mL; Corning, 354233) or Geltrex-coated plates in MEF-CM with 10 μM Y-27632 at a density of 10,000 cells/cm². On the following day, cells were cultured in 5% CO₂ and 5% O₂ and medium was changed to TE differentiation medium: DMEM/F12, with 1× ITS-X (Gibco, 41400045), 64 μg/mL L-ascorbic acid (Sigma, A4544), 543 μg/mL NaHCO₃ (Sigma, S5761), 2% BSA (Gibco, 15260037), 10 ng/mL BMP4, and 2 μM IWP2 (STEMCELL Technologies, 72124). The medium was refreshed daily for 4 days. After 4 days, the medium was switched to modified TSC medium: Advanced DMEM/F12 (Gibco, 12634010) supplemented with 1× N2,1× B27, 1×L-glutamine, 0.1 mM β-mercaptoethanol, 0.05% BSA, 1% knockout serum replacement (KSR; Gibco, 10828028), 2 μM CHIR99021 (STEMCELL Technologies, 72054), 0.5 μM A83-01, 1 μM SB431542 (STEMCELL Technologies, 72232), 5 μM Y-27632, 0.8 μM VPA, 100 ng/mL basic FGF, 50 ng/mL recombinant human EGF (Peprotech, AF-100-15), 50 ng/mL recombinant human HGF (PeproTech, 100-39), and 20 ng/mL Noggin (R&D Systems, 6057-NG). The medium was refreshed every other day.

### Derivation of TSC from primed hPSC through chemical conversion (ccTSC)

Derivation of ccTSC was preformed according to a previous chemical induction protocol with modifications^25^. In brief, hPSCs were dissociated using Accutase and plated onto collagen IV (5 μg/mL), Geltrex-coated plates with 10 μM Y-27632 at a density of 10,000 cells/cm². Two days after plating, the medium was changed to N2B27 medium supplemented with 10 ng/mL LIF, 1 μM PD0325901, and 1 μM VPA and cells were cultured in 5% CO₂ and 5% O₂. After three days in this medium, it was changed to TSC medium: DMEM/F12 supplemented with 0.1 mM β-mercaptoethanol, 0.2% FBS (Sigma, F4135), 0.5% Penicillin-Streptomycin (P/S; Gibco, 15140122), 0.3% BSA, 1% ITS-X (Gibco, 41400045), 1.5 μg/mL L-ascorbic acid, 50 ng/mL EGF, 2 μM CHIR99021, 0.5 μM A83-01, 1 μM SB431542, 0.8 mM VPA, and 5 μM Y-27632. The medium was refreshed every other day.

### Derivation of TSCs from naïve hPSCs (nTSC)

Naïve hPSCs were dissociated using Accutase and plated onto collagen IV (5 μg/mL), Geltrex-coated, or MEF plates in PXGL medium supplemented with 10 μM Y-27632 and 1% Geltrex at a density of 50,000 cells/cm². Cells were cultured in 5% CO₂ and 5% O₂. After two days, the medium was changed to TSC medium.

### Derivation of TSCs from the naïve-hPSC-derived blastoids (nbTSC)

Blastoids transferred on IBIDI 8 wells coated with laminin-521 (10 μg/mL; BioLamina, LN521) in IVC1 medium^84^, composed of 50% DMEM/F12, 50% Neurobasal, 20% FBS, 1× L-glutamine, 1× ITS-X, 8 nM β-estradiol (Sigma, E2758), 200 ng/mL progesterone (Sigma, P8783), and 25 μM N-acetyl-L-cysteine (Sigma, A9165). Blastoids were cultured in 5% CO₂ and 5% O₂. Two days post-attachment, the medium was switched to TSC medium. Within 4-5 days, outgrowths were observed. After 10 days of attachment, the outgrowths were dissociated with Accutase and passaged onto either collagen IV (5 μg/mL) coated or MEF plates.

### Maintenance of TSCs

TSCs were passaged using TrypLE™ Express (Gibco, 12605010) when 80% confluent (approximately every 4 days) at a ratio of 1:4-6 onto collagen IV (5 μg/mL) or Geltrex-coated plates. G-banding karyotyping of cells from passage 10-20 was conducted at Cell Line Genetics to verify the chromosomal status of each TSC line. Cells from passage 6-15 were used for downstream differentiation experiments unless specified otherwise.

### Differentiation of TSCs

TSCs were grown until 80% confluent, then dissociated with TrypLE™ Express for 10-15 minutes at 37°C.

For the induction of EVTs, TSCs were seeded into 6-well plates pre-coated with Geltrex or 1 μg/mL collagen IV at a density of 0.75 × 10⁵ cells per well and cultured in EVT1 medium supplemented with 2% Geltrex. On day 3, the medium was replaced with EVT2 medium supplemented with 0.5% Geltrex. Cells were analyzed on day 6. Alternatively, the cells were further dissociated and passaged onto a new collagen IV (1 μg/mL) plate with EVT2 medium (without KSR) supplemented with 1% Geltrex for an additional 2 days.

For the induction of STs, TSCs were seeded into 6-well plates pre-coated with Geltrex or 3 μg/mL collagen IV at a density of 1 × 10⁵ cells per well and cultured in ST medium, consisting of DMEM/F12, 0.1 mM β-mercaptoethanol, 0.5% P/S, 0.3% BSA, 1% ITS-X, 2.5 μM forskolin (Sigma, F6886), 2.5 μM Y-27632, and 4% KSR. The medium was replaced on day 3, and cells were analyzed on day 6.

### Trophoblast organoid

TOs were generated following previously established protocols^46,48^. Briefly, TSCs were grown until they confluent, then dissociated with TrypLE™ Express and washed twice. A total of 3,000 cells were suspended in 30 μL Matrigel droplets (Corning, 354277), seeded into 24-well plates and incubated at 37°C for 30 minutes before adding trophoblast organoid medium (TOM), which consisted of Advanced DMEM/F12, 1× N2 supplement, 1× B27 supplement, 1.25 mM N-acetyl-L-cysteine, 2 mM L-glutamine, 50 ng/mL recombinant human EGF, 1.5 μM CHIR99021, 80 ng/mL recombinant human R-spondin-1 (R&D Systems, 4645-RS), 100 ng/mL recombinant human FGF-2, 50 ng/mL recombinant human HGF, 500 nM A83-01, 2.5 μM prostaglandin E2 (Sigma, P0409), 0.1 mM 2-mercaptoethanol, and 2 μM Y-27632. Medium was refreshed every other day. Organoids were maintained for 8-10 days between passages. For passage, Matrigel droplets were broken up in cold DMEM/F12, and organoid pellets were dissociated with TrypLE™ Express for 20 minutes after centrifugation. After three washes with DMEM/F12, cells were passed through a 70 μm strainer to remove large and unbroken chunks, and 3,000-5,000 cells were re-seeded in 30 μL Matrigel droplets.

EVT-TOs were generated following a previous protocol with minor modification^49^. Briefly, TOs were cultured in TOM for 3-5 days after seeding or until reaching a size of 100 μm, then switched to EVT1 medium, consisting of DMEM/F12, 0.1 mM β-mercaptoethanol, 0.5% P/S, 0.3% BSA, 1% ITS-X, 100 ng/mL NRG1, 7.5 μM A83-01, 2.5 μM Y-27632, and 4% KSR. After 5 days in EVT1 medium, it was replaced with EVT2 medium (EVT1 medium minus NRG1) for an additional 3 days. All organoid media were refreshed every 2-3 days.

### Proliferation rates of TSCs and trophoblast organoids

The confluency of TSCs was assessed using CELLCYTE Studio (CYTENA) to scan live cell development. Confluency area measurements were used to analyze proliferation rates within 48 hours of scanning. The regions of interest (ROIs) of trophoblast organoids were analyzed in ImageJ via the polygon selection tools. Measurements of the ROIs included area (in square pixels) and perimeter (in pixels). Area values were normalized to the mean area at the initial time point for each condition. Means and standard errors were calculated for each condition at three time points: day 2, day 4, and day 7.

### Fluorescence in situ hybridization and chromosome number measurement

TSCs were grown until confluent, dissociated with TrypLE™ Express, and incubated in hypotonic solution (0.075 M KCl, Sigma, P5405) at 37°C for 17 minutes. Cells were fixed with Carnoy’s fixative (3:1 methanol [Sigma, 494437] to acetic acid [Sigma, A6283]) and washed three times. A suspension at a concentration of 3 million cells/mL was dropped onto slides and air-dried at room temperature. Slides were then washed in 2× SSC buffer (Invitrogen, 15557044) for 2 minutes, followed by sequential ethanol washes (70%, 85%, 100%), 2 minutes each. The combined satellite Enumeration Probes for chromosomes 7, 12, and 18 (3 μL each, Oxford Gene Technology) and hybridization solution (1 uL, Oxford Gene Technology) were added, 22 mm × 22 mm coverslips were applied, and slides were denatured at 75°C for 2 minutes before overnight incubation at 37°C. The next day, slides were washed in 0.4× SSC preheated to 72°C for 2 minutes, then incubated in 2× SSC with 50 μg/mL DAPI for 10 minutes.

Imaging was conducted at 40× magnification, with z-stacks acquired at a minimum interval of 7 μm. Maximum intensity projections were generated using ImageJ to identify and count nuclear puncta for each probe. As cell cycle synchronization was not performed, cell cycle stages were carefully assessed, with S phase cells identified based on closely spaced centromeric signals for any chromosome. In S phase cells, only widely separated puncta for the same chromosome were considered distinct copies. Cells with unreliable counts for two or more probes were excluded from the analysis. Cells showing ≥4 signals for at least two probes were classified as tetraploid (4n). Chromosomes with signal counts deviating from the whole-cell ploidy were categorized as aneuploid. Average of 300, minimum of 100 cells were analyzed for aneuploidy across multiple imaging fields.

### Immunofluorescence staining

Cells or organoids were fixed with 4% paraformaldehyde (PFA, Electron Microscopy Sciences, 15710) for 30 minutes. After three washes with PBS for 5 minutes each, samples were blocked and permeabilized with blocking buffer (PBS supplemented with 3% normal donkey serum [Sigma, S30] and 0.3% Triton X-100 [Sigma, T8787]) for 30 minutes. Cells were then incubated with primary antibodies diluted in blocking buffer for 1 hour, followed by three PBS washes. Secondary antibodies conjugated to Alexa Fluor and 10 ng/mL DAPI were added for 30 minutes, followed by three PBS washes. All antibodies used are listed in the Table S3.

### RNA isolation and Real-time qPCR

RNA was extracted using the RNeasy Mini Kit (Qiagen, 74106) following the manufacturer’s protocol. cDNA was synthesized using iScript™ Reverse Transcription Supermix (Bio-Rad, 1708840). Quantitative RT-PCR was performed using PowerUp™ SYBR™ Green Master Mix (Applied Biosystems, A25742) on the 7900HT Fast Real-Time PCR System (Applied Biosystems). The cDNA was used as the RT-PCR template, and gene expression was analyzed using the delta-delta cycle threshold method with GAPDH as a housekeeping gene. Error bars represent the standard deviation of biological replicate fold-change values. All primers used for RT-PCR are listed in Table S4.

### Bulk RNA sequencing

Total RNA was isolated using the RNeasy mini Kit. The purity and concentration of the RNA was assessed using a NanoDrop 2000 spectrophotometer. Library preparation was performed using the Optimal Dual-mode mRNA Library Prep Kit (BGI-Shenzhen, China). The double-stranded cDNA product was converted to blunt ends through end repair reactions. Library products were amplified via PCR and subjected to quality control. Single-stranded library products were generated through denaturation, followed by a circularization reaction to produce single-stranded circular DNA products. Sequenced reads of 100 bases in length were generated on the G400 platform. Raw sequencing data was processed with SOAPnuke^85^ to remove low quality reads or reads with adaptor sequence. Clean sequencing reads were mapped to human reference genome (hg38) with Bowtie2^86^ and gene expression quantification was performed with RSEM^87^ as read counts. CPM (counts per million reads) values were used as gene expression levels. Differential expression analysis was run by Deseq2^88^. Heatmaps, principal component analyses (PCA), scatter plot, venn diagram, volcano plot, KEGG pathway enrichment, Gene Ontology analysis were performed using R software (version 4.3.3).

### Whole genome bisulfite sequencing

DNA was extracted using Qiagen DNeasy Blood & Tissue Kit. Library preparation was performed using the MGIEasy Whole Genome Bisulfite Sequencing Library Prep Kit (BGI-Shenzhen, China). For library construction, genomic DNA was mixed with 1% unmethylated lambda phage DNA and then sheared into small fragments using the Covaris LE220 Focused-ultrasonicator. DNA fragments were treated with bisulfite using the EZ DNA Methylation Gold Kit (Zymo Research, D5005). Bisulfite-treated library products were amplified through PCR and subjected to quality control. Sequenced reads of 150 bases in length were generated on the G400 platform. Raw sequencing data were processed with SOAPnuke to remove low-quality reads or reads containing adapter sequences^89^. Clean sequencing reads were mapped to the GRCh38 reference genome using Bismark^90^. DNA methylation levels were calculated as described in a previous publication^91^. Promoters were defined as regions 1 kb upstream and downstream of transcription start sites. CpG methylation levels with ≥5 reads were analyzed.

For copy number variance analysis, chromosomes were divided into 1 Mb windows, and RPM (reads per million mapped reads) was calculated for each window. These values were normalized against karyotypically normal hPSCs and smoothed across 10 Mb intervals. Only windows with over 50 RPM in control hPSCs were analyzed. Copy number gains and losses were identified using thresholds of 2.3 and 1.7 copies, respectively^92^.

### Chromosomal Analysis from snATAC-seq Data

Placental snATAC-seq data were obtained from the public accession GSE247038^71^. Cell Ranger atac-count was run on the FASTQ files for each donor to obtain fragments.tsv files, which were then analyzed by epiAneufinder^76^ to determine individual nuclei copy number changes in the software R^93^. To classify a cell as aneuploid, 60% or more of the bins on any chromosome had to be labeled as a CNV by epiAneufinder. Non-nuclei were filtered out by modifying the minFrags parameter based on quality control metrics from Cell Ranger’s output files, while all other parameters were set to default. Chromosomes X, Y, and mitochondrial DNA were excluded from analysis, with UCSC hg38 used as the reference genome. Human PBMC ATAC-seq data were obtained from 10x Genomics’ publicly available datasets. The log10 fragments of nuclei for each donor and the PBMC controls were calculated using custom scripts in R and visualized using ggplot2^94^.

To quantify and compare aneuploidy rates across groups (“early placenta”, “term placenta”, or “PBMC”), a binomial generalized linear mixed model (GLMM) was used. The model was implemented with the R package lme4^95^, where the response variable was a binary indicator denoting whether a given cell was inferred to be aneuploid based on epiAneufinder’s results. Cell type was included as a fixed effect factor predictor, while tissue donor was included as a random effect grouping factor to account for non-independence among nuclei from the same donor. The fitted model was interpreted using the R package “marginaleffects”^96^ to compute predicted values and comparisons of aneuploidy rates for each cell type, averaging over levels of the random effect term (tissue donor). Results were visualized using ggplot2^94^ and ggsignif^97^ packages in R.

### Chromosomal Analysis from scRNA-seq Data

Placental SmartSeq2 RNA data were obtained from the public accession GSE89497^70^. Pooled FASTQ files were split into individual cells using code provided by the authors of the accession and aligned to UCSC hg38 using Salmon^98^ with default settings. The resulting count matrix was aggregated to the gene level using the R package “tximport”^99^ and cell identities were determined through UMAP plotting and marker gene expression using Seurat^100^.

The R package “scploid”^72^ was used to determine aneuploidy, with a min.median value of 19 chosen to minimize residuals while achieving acceptable criteria for all three of the package’s QC metrics (zeros, residuals, and genes). All other parameters were set to default. Only clusters with EVT or CTB identities that received acceptable scores on all three QC metrics were considered for analysis.

### Autophagy inhibition

Chloroquine (Sigma, C6628) was used to inhibit autophagy in TSCs. Briefly, TSCs and parental primed hPSCs were seeded on Geltrex-coated 12-well plates at a density of 5 × 10⁴ cells/well. Cells were treated with 0, 10 μM, and 50 μM chloroquine. The culture media included TSC medium for TSCs and mTeSR medium with Y-27632 for hPSCs. Cells were collected at 24 and 48 hours for downstream analysis of apoptotic rates using flow cytometry.

### Flow cytometry

Cells were dissociated into single cells using TrypLE™ Express and passed through a 40 μm strainer.

For cell cycle analysis, cells were fixed with 4%PFA for 15 minutes at room temperature. After two washes in PBS, cells were incubated with (concentration) DAPI for 15 minutes at room temperature. Cells were then washed twice in PBS and resuspended in PBS supplemented 2% fetal bovine serum (FBS) for flow cytometry analysis.

For studying autophagy inhibition, all supernatant apoptotic cells were collected along with dissociated single cells, passed through a 40 μm strainer, and stained with SYTOX™ Green (1:100, Invitrogen, S7020), a marker specific for apoptotic cells, for 20 minutes at room temperature. After washing with PBS supplemented with 2% FBS, cells were fixed with 4% PFA for 15 minutes at room temperature. Following two washes in PBS, cells were resuspended in 2% FBS/PBS for flow cytometry analysis.

Flow cytometry was performed using a FACSCanto II (BD Biosciences), and results were analyzed using FlowJo software.

### Western Blot Analysis

Protein from cultured cells was harvested using NP-40 lysis buffer (Boston Bioproducts, BP-119x) with added protease inhibitors (Santa Cruz, sc-24948). Protein concentrations were determined using the BCA Protein Assay (Thermo Scientific, 23225). A total of 25 μg of protein from each lysate was subjected to electrophoresis on a 4-12% Bis-Tris gel (Invitrogen, NW04120BOX) and transferred onto a PVDF membrane (Life Technologies, LC2002). The membrane was then sectioned based on the molecular weights of the markers being analyzed. The membrane was blocked for 1 hour at room temperature in 5% nonfat milk in TBS-T (Tris-buffered saline [sigma,T5912] with 0.1% Tween-20 [Thermo Scientific, J20605.AP]) and incubated overnight at 4°C with primary antibodies while gently rocking. Membranes were rinsed in TBS-T and treated with a species-specific horseradish peroxidase-conjugated secondary antibody for 1 hour at room temperature. All antibodies used are listed in the Table S3. Following additional washes with TBS-T, the membranes were developed to visualize the bound antibodies using SuperSignal West Pico Plus Chemiluminescent Substrate (Thermo Scientific, 34580).

## Statistical analysis

Unless otherwise specified, results are expressed as the mean ± standard deviation (SD). Statistical analyses were performed with at least three biological replicates, unless stated otherwise. Statistical significance was determined using unpaired ANOVA and student’s t-test performed with GraphPad Prism 10 or R software (version 4.3.3). A p-value < 0.05 was considered statistically significant.

## Ethics Statement

This study was reviewed and approved by the Embryonic Stem Cell Research Oversight (ESCRO) Committee at the University of Washington.

## Acknowledgments

We thank all members of the Stem Cell Genomics team, including the Yang and Hamazaki Laboratories, for their helpful feedback and discussions. We are grateful to Dr. Ali Brivanlou for providing RUES2 cells, as well as for his mentorship and support during M.Y.’s smooth transition to establishing her independent lab. We appreciate Dr. Lucia Vjotech and Dr. Shahrokh Paktinat for their assistance in setting up western blotting protocols. We thank Dr. Dale Hailey from the Garvey Imaging Core at UW for confocal imaging support, and Thane Mittelstaedt and Delphina Walker-Phelan from the UW Cell Analysis Facility for their help with flow cytometry. We are grateful to Erica Jonlin, ISCRM Regulatory Manager, for her guidance on the ESCRO application process. We also thank Olivia Waters from the Yang Lab for assisting with FISH slide imaging and Dr. Jenny Morse from OGT for her assistance in establishing the FISH protocol. We thank Dr. Hao Ming and Dr. Zongliang Jiang for their feedback on WGBS analysis. This work was supported by the NICHD Path to Independence Award (4R00HD107219-03), startup funding from UW department of OB/GYN, ISCRM, and the Brotman Baty Institute (BBI).

## Author contributions

M.Y. conceptualized the study. Experimental design was carried out by M.Y. and D.W., under the supervision of M.Y. Cell culture experiments, microscopy, and data analysis were performed by D.W. and M.Y. Bulk sequencing data analysis was conducted by D.W. FISH and quantification were performed by K.L. Placental single-cell analysis was carried out by A.C. and R.M. TO organoid image quantification was conducted I.J.

**Table S1** Karyotypes of TSCs.

**Table S2** Quality Control of Placental scRNA-seq Data for scPloidy Analysis

**Table S3** Antibodies used in this study.

**Table S4** qRT-PCR primers.

**Figure S1.**
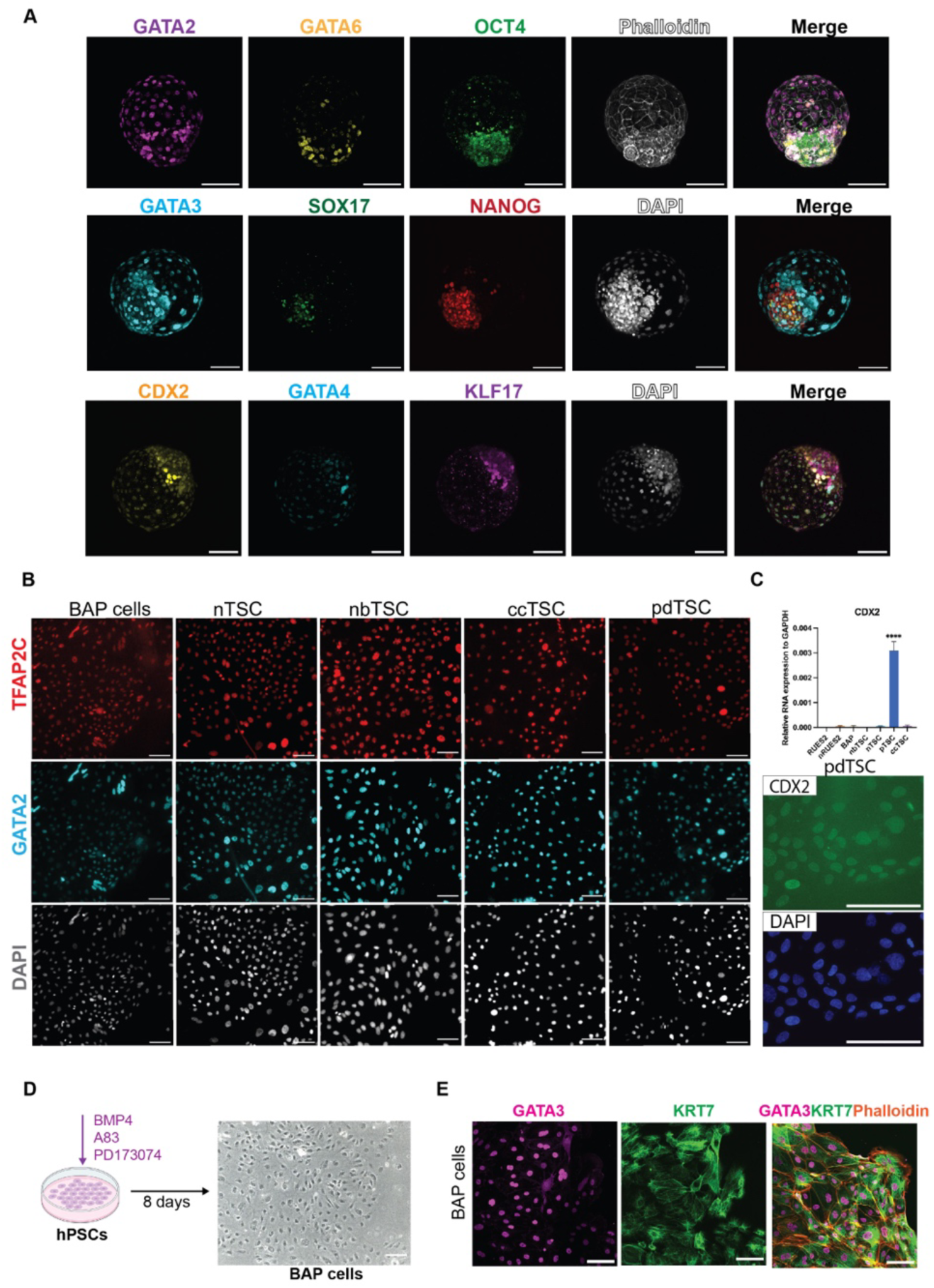
Validation of TSC derivation and BAP cells (related to Figure 1) (A) Immunostaining of trophoblast markers (GATA2, GATA3), trophectoderm marker (CDX2), extraembryonic endoderm markers (GATA6, SOX17, GATA4), and pluripotency markers (OCT4, NANOG, KLF17) in blastoids derived from naïve hPSCs. Scale bar, 100 μm. (B) Immunostaining of additional trophoblast markers, TFAP2C and GATA2, in BAP cells, nTSCs, nbTSCs, ccTSCs, and pdTSCs. Scale bar, 100 μm. (C) Relative expression of CDX2 showing significantly elevated levels in pdTSCs, with no detectable expression in other TSCs or hPSCs. \**P* < 0.05. Immunostaining confirms CDX2 expression in pdTSCs. Scale bar, 100 μm. (D) Schematic representation of BAP cell derivation from hPSCs. Scale bar, 100 μm. (E) Immunostaining of trophoblast markers GATA3 and KRT7 in BAP cells. Scale bar, 100 μm.

**Figure S2.**
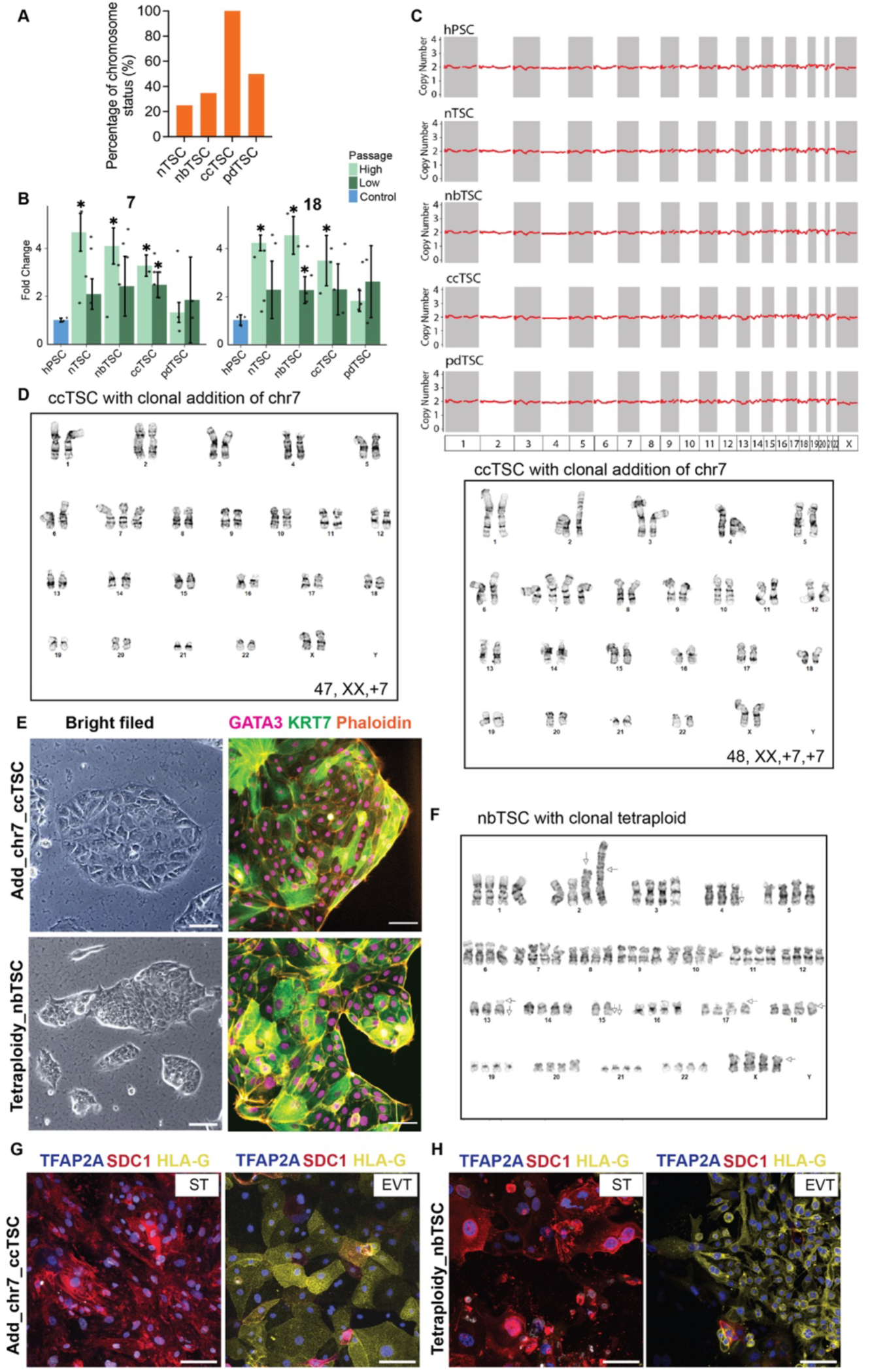
Detection and validation of aneuploid TSCs (related to Figure 2) (A) Percentage of cells with abnormal karyotypes detected in TSC lines from G-banding karyotyping results. 20-40 cells were assessed for each TSC type. (B) Fold change in percentage of aneusomic cells per chromosome for chromosomes 7 and 18 in TSCs at passage 3, normalized to the baseline control primed hPSCs. \**P* < 0.05 compared to primed hPSCs. (C) Chromosome analysis of TSCs through WGBS copy number variance detection. (D) Representative G-banding karyotype of the ccTSC line showing a clonal effect with an additional copy of chromosome 7. (E) Bright-Tield images and immunostaining of trophoblast markers GATA3 and KRT7 in ccTSCs and nbTSCs with clonal aneuploidy. Scale bar, 100 μm. (F) Representative G-banding karyotype of the nbTSC line showing a clonal effect of tetraploidy. (G) and (H) ST and EVT differentiation validation in ccTSCs and nbTSCs with clonal aneuploidy. Immunostaining for ST marker SDC1, EVT marker HLA-G, and general trophoblast marker TFAP2A. Scale bar, 100 μm. Immunostaining of SDC1 in TSC_EVTs showing spontaneous differentiation of STs. Scale bar, 100 μm.

**Figure S3.**
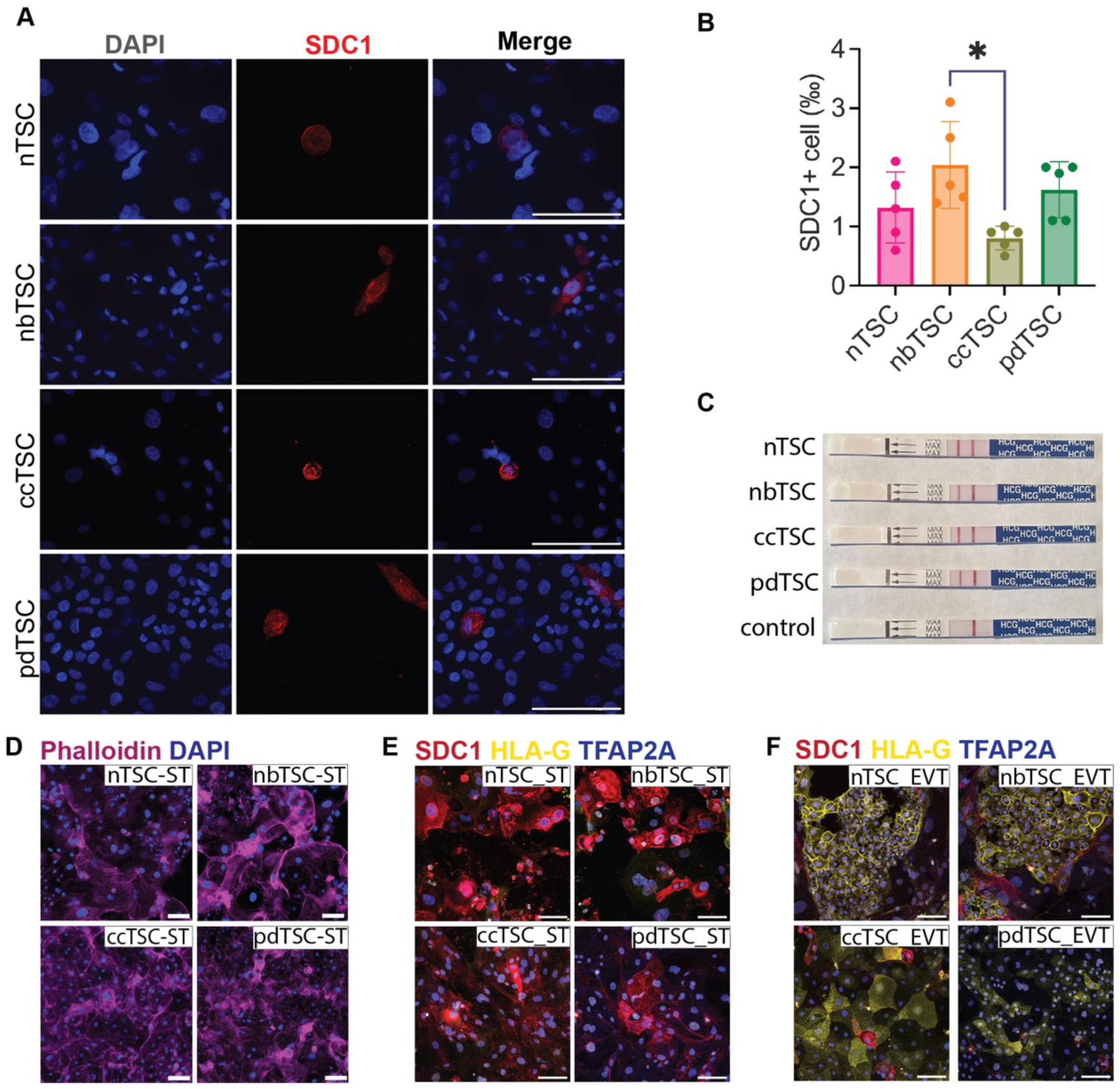
Differentiation capacity of TSCs is uncompromised by CIN (related to figure 3) (A) Immunostaining of SDC1 in TSCs showing spontaneous differentiation of STs. Scale bar, 100 μm. (B) Permillage of spontaneous STs in TSCs. *P < 0.05. (C) HCG test (OTC pregnancy test) results in TSC culture medium compared to medium control, showing secretion of HCG by spontaneously differentiated STs. (D) Phalloidin and DAPI staining for cytoskeleton and nuclear visualization, respectively, demonstrating cell fusion in STs. Scale bar, 100 μm. (E) Validation of ST differentiation from TSCs with minimal aneuploidy (passage 3). Immunostaining for ST marker SDC1, EVT marker HLA-G, and general trophoblast marker TFAP2A. Scale bar, 100 μm. (F) Validation of EVT differentiation from TSCs with minimal aneuploidy (passage 3). Immunostaining for ST marker SDC1, EVT marker HLA-G, and general trophoblast marker TFAP2A. Scale bar, 100 μm.

**Figure S4.**
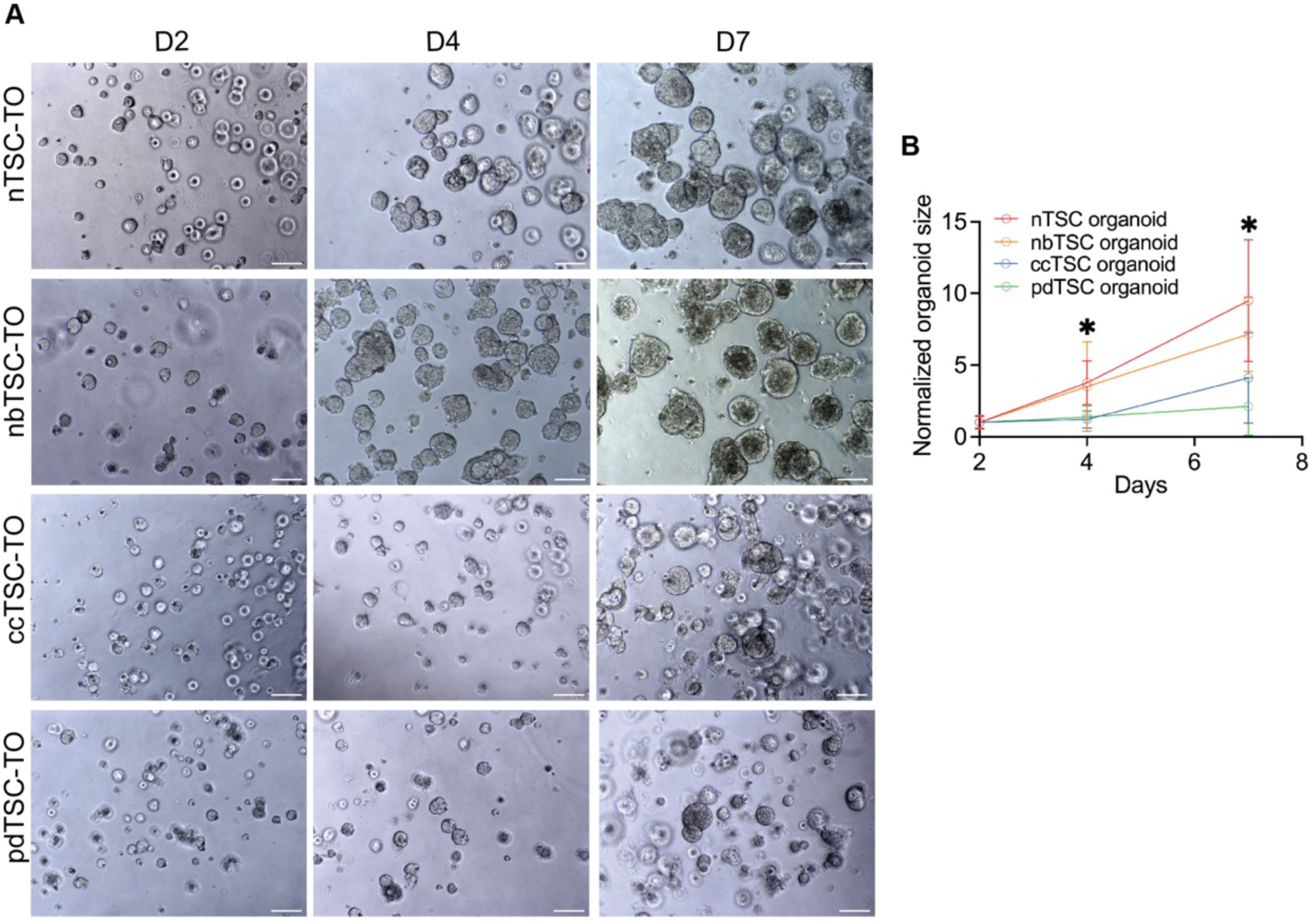
Trophoblast Organoid Growth of TSCs. (A) Bright-Tield images of trophoblast organoids (TO) derived from nTSCs, nbTSCs, ccTSCs, and pdTSCs at days 2, 4, and 7 post-seeding. Scale bar, 100 μm. (B) Normalized size of trophoblast organoids at days 2, 4, and 7. \**P* < 0.05. At Day 4, nTSC vs ccTSC, nTSC vs pdTSC, nbTSC vs ccTSC, nbTSC vs pdTSC showing signiTicant difference. Same pattern showing at Day 7.

**Figure S5.**
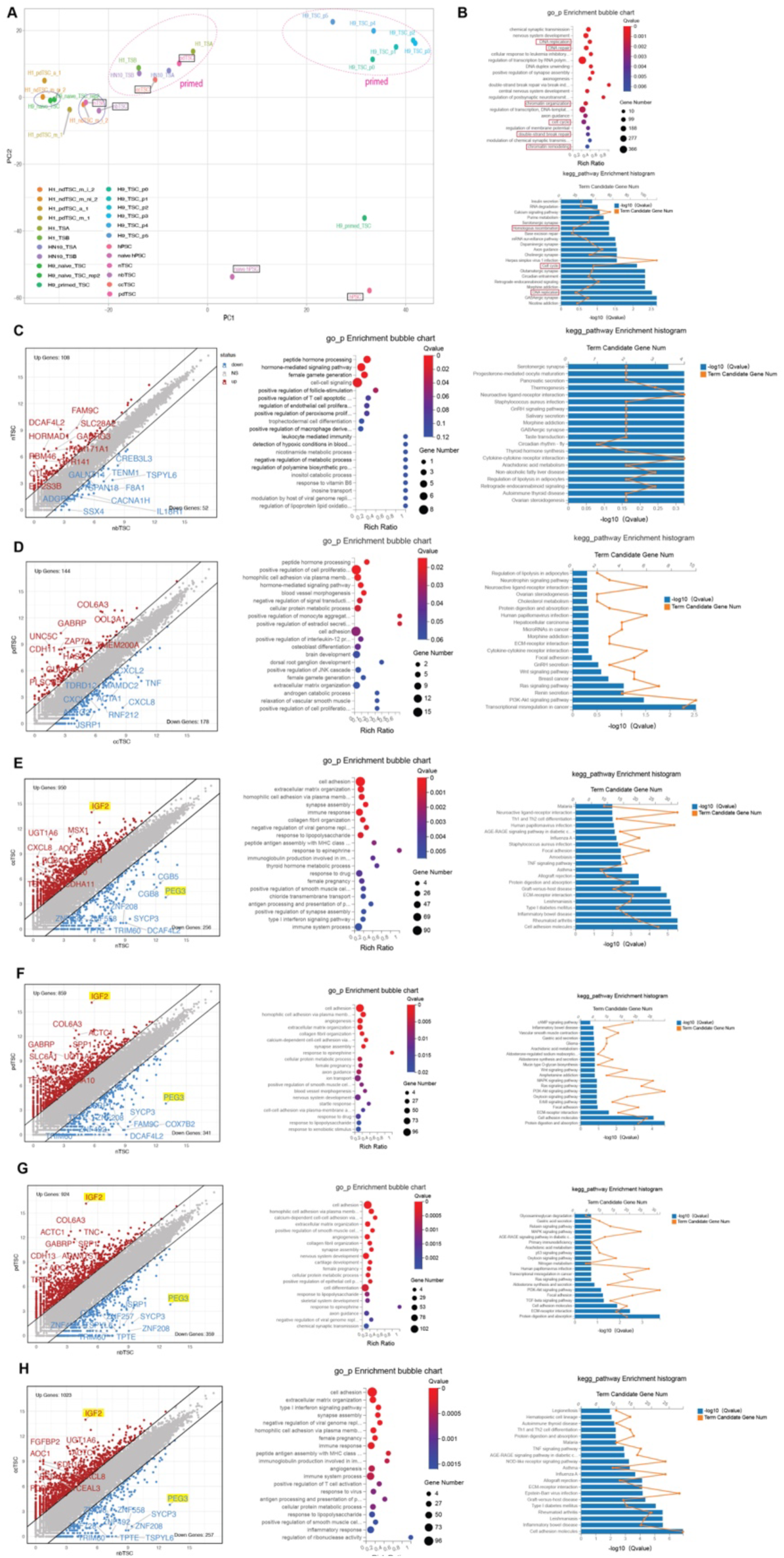
Transcriptome analysis of TSCs. (A) PCA of all TSCs derived in this manuscript and the parental hPSCs (within black rectangle) alongside other previously published TSC datasets, including: **H1_ndTSC_m_i_2**, **H1_ndTSC_m_ni_2**, H1_pdTSC_a1, H1_pdTSC_m1^23^; H1_TSA, H1_TSB, HN10_TSA, HN10_TSB^19^; **H9_naive_TSC**, **H9_naive_TSC-rep2**, H9_primed_TSC^13^; H9_TSC_p0, H9_TSC_p1, H9_TSC_p2, H9_TSC_p3, H9_TSC_p4, H9_TSC_p5^22^. Text in bold indicates naıïve TSCs, while the remaining entries represent primed TSCs. (B) GO biological process and KEGG pathway enrichment analysis of downregulated genes in TSCs compared to primed hPSCs, showing signiTicant enrichment in DNA replication and repair, chromatin organization and remodeling, cell cycle, double-strand break repair, and homologous recombination. (C) and (H) Left panel: scatter plots showing pairwise transcriptome comparisons between each TSC type. Middle and right panels: GO biological process and KEGG pathway enrichment analyses for upregulated genes in each pairwise comparison.

**Figure S6.**
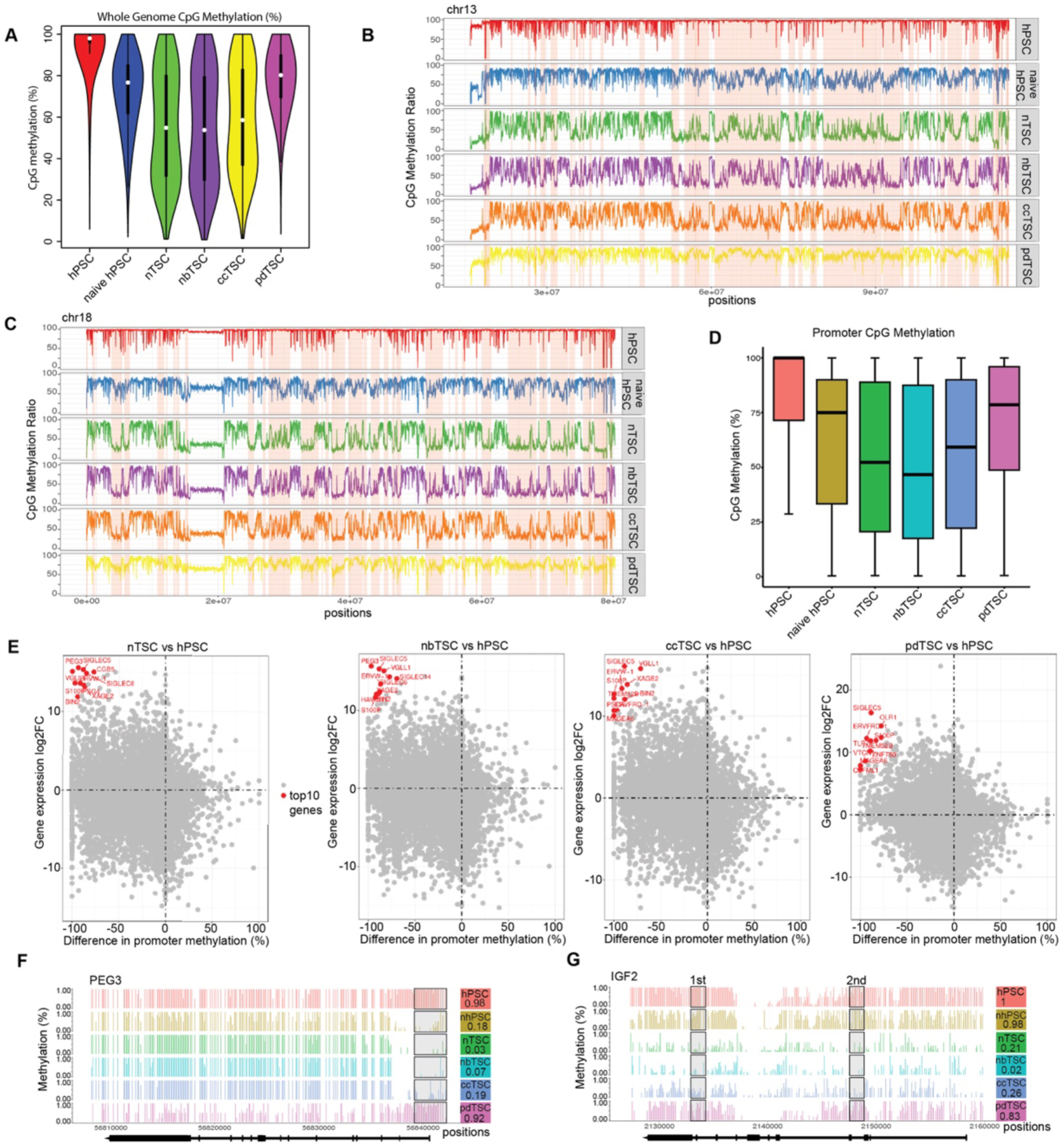
DNA methylome profiling of TSCs. (A) Violin plots of whole genome CpG methylation levels in nTSCs, nbTSCs, ccTSCs, pdTSC, and parental hPSCs. White dots indicated the median, and boxplot lines are overlaid within the violin plots. (B) DNA methylation patterns across chromosome 13 in TSCs and parental hPSCs. The vertical axis represents the CpG methylation ratio, with PMDs highlighted in pink. (C) DNA methylation patterns across chromosome 18 in TSCs and parental hPSCs. The vertical axis represents the CpG methylation ratio, with PMDs highlighted in pink. (D) Boxplot of promoter CpG methylation levels in nTSCs, nbTSCs, ccTSCs, pdTSC, and parental hPSCs. (E) Scatter plot of log2 fold change (log2 FC) of gene expression from transcriptome data versus the difference in promoter methylation levels between TSCs and primed hPSCs. The top 10 genes are labeled in red. (F) DNA methylation patterns at the PEG3 locus. The promoter region is within rectangle, with corresponding methylation levels indicated on the right. (G) DNA methylation patterns at the IGF2 locus. The IGF2 DMRs are within rectangle, with corresponding methylation levels indicated on the right. DMR0 (GRCh38: 11,2148236-2148307) and DMR1 (GRCh38: 11, 2133452-2133743).

**Figure S7.**
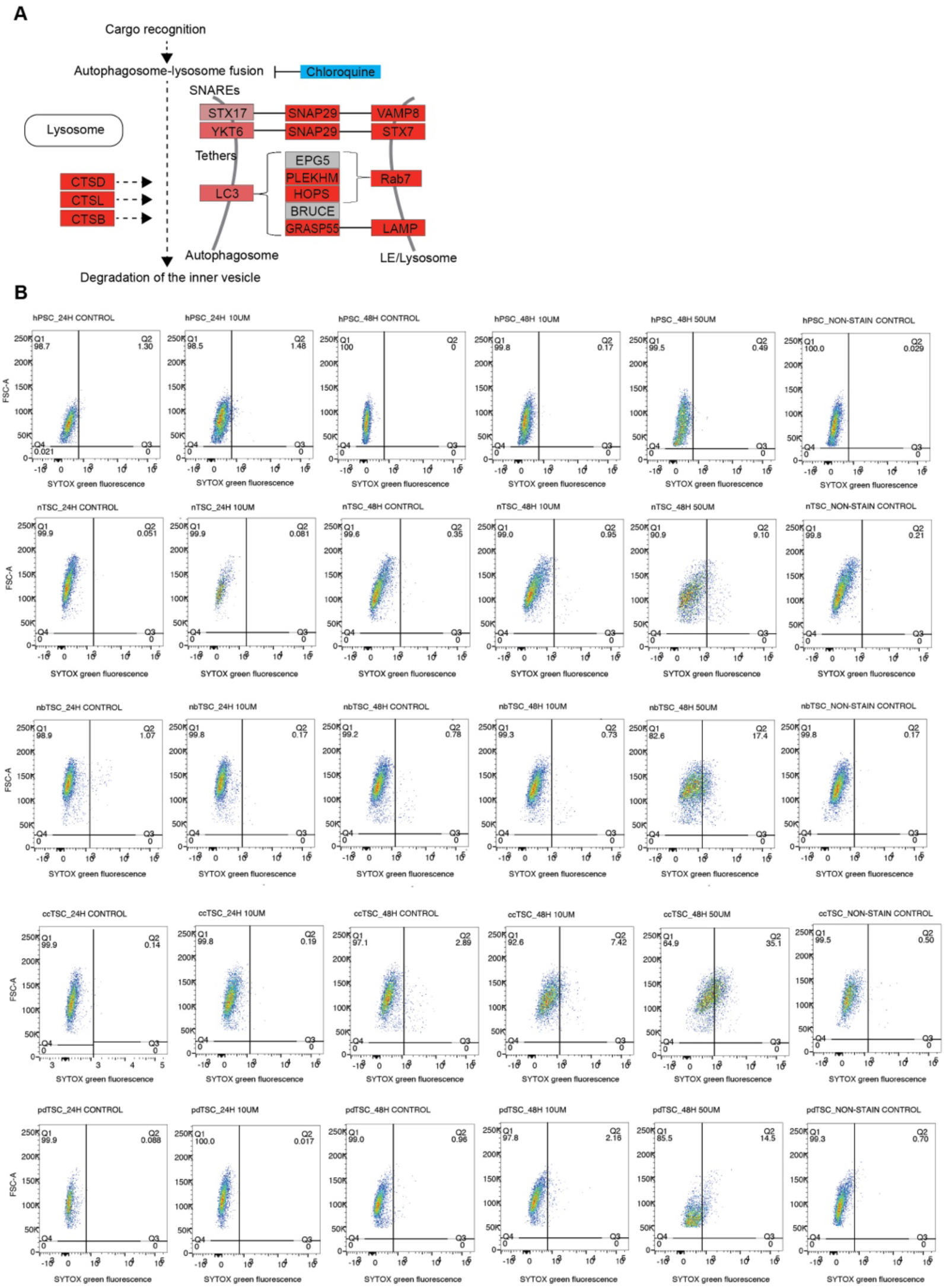
Chloroquine Inhibition of Autophagosome-Lysosome Fusion Increases Cell Death in TSCs. (A) Transcriptome comparison indicates upregulation (highlighted in red) of genes associated with the autophagosome-lysosome fusion (KEGG pathway visualization). CQ treatment is used to inhibit autophagosome-lysosome fusion. (B) Flow cytometry analysis of cell viability using Sytox Green staining in TSCs and hPSCs following CQ treatment (10 µM and 50 µM) for 24 and 48 hours. TSCs show greater sensitivity to CQ treatment compared to hPSCs, indicating increased cell death.

**Figure S8.**
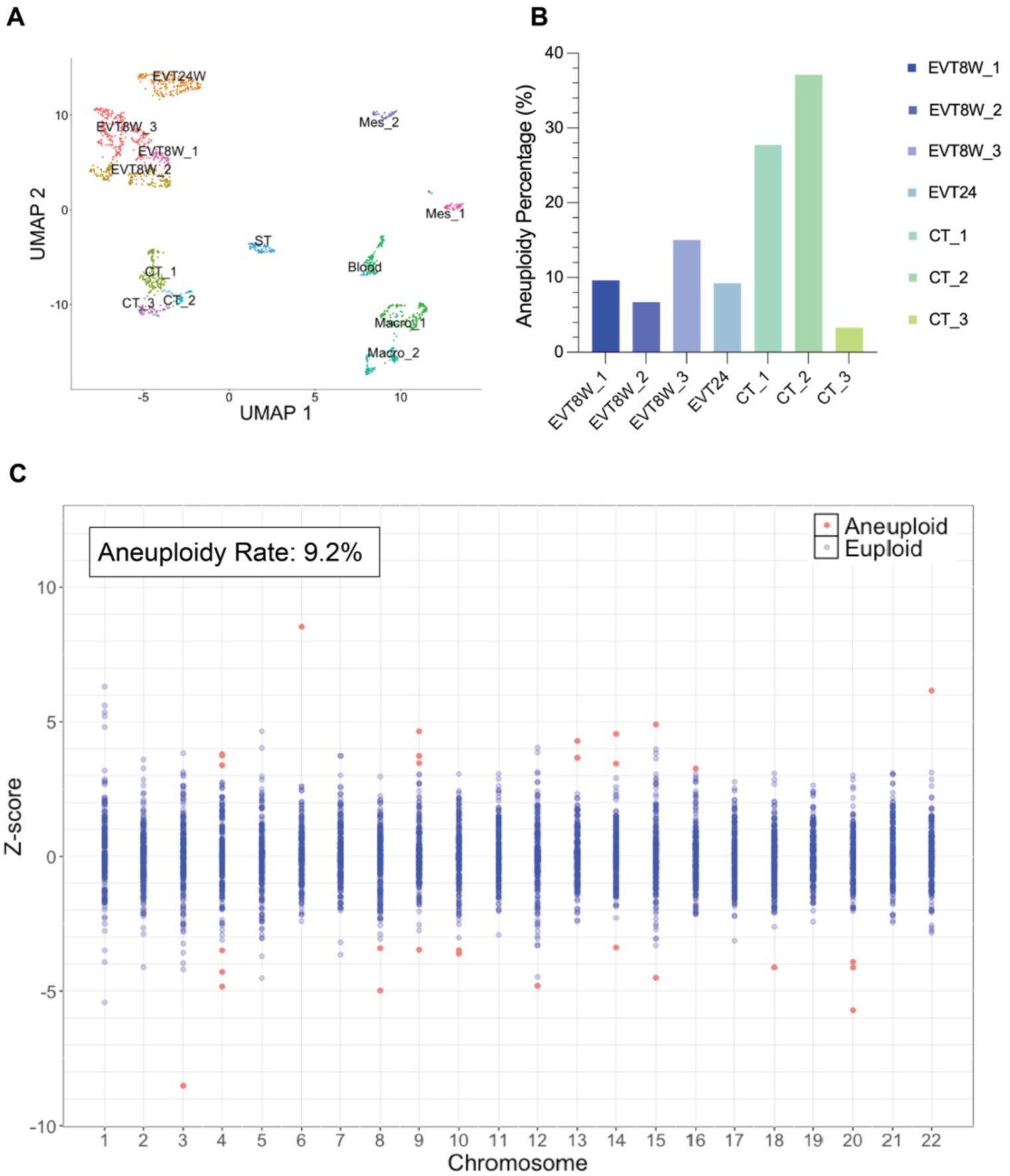
Aneuploidy Detection in Developing Human Placentas Using scRNA-seq (Smart2-seq) (A) UMAP of scRNA-seq data from Tirst trimester (8 weeks) and second trimester (24 weeks) placentas, generated using Seurat. Cell type identities for each cluster were determined by marker gene expression. Mes, mesenchyme; Macro, macrophage. (B) Percentage of aneuploid cells in EVT and CT Seurat clusters. Aneuploidy calling was conducted within the clusters using the SCPloid package in R. (C) Distribution of z-score values for each chromosome per cell in 24-week EVT cluster (EVT24W). Red dots indicate chromosomes with copy number changes as identiTied by SCPloid. The percentage of putative aneuploid cells in 24-week EVT is 9.2%.

**Figure S9.**
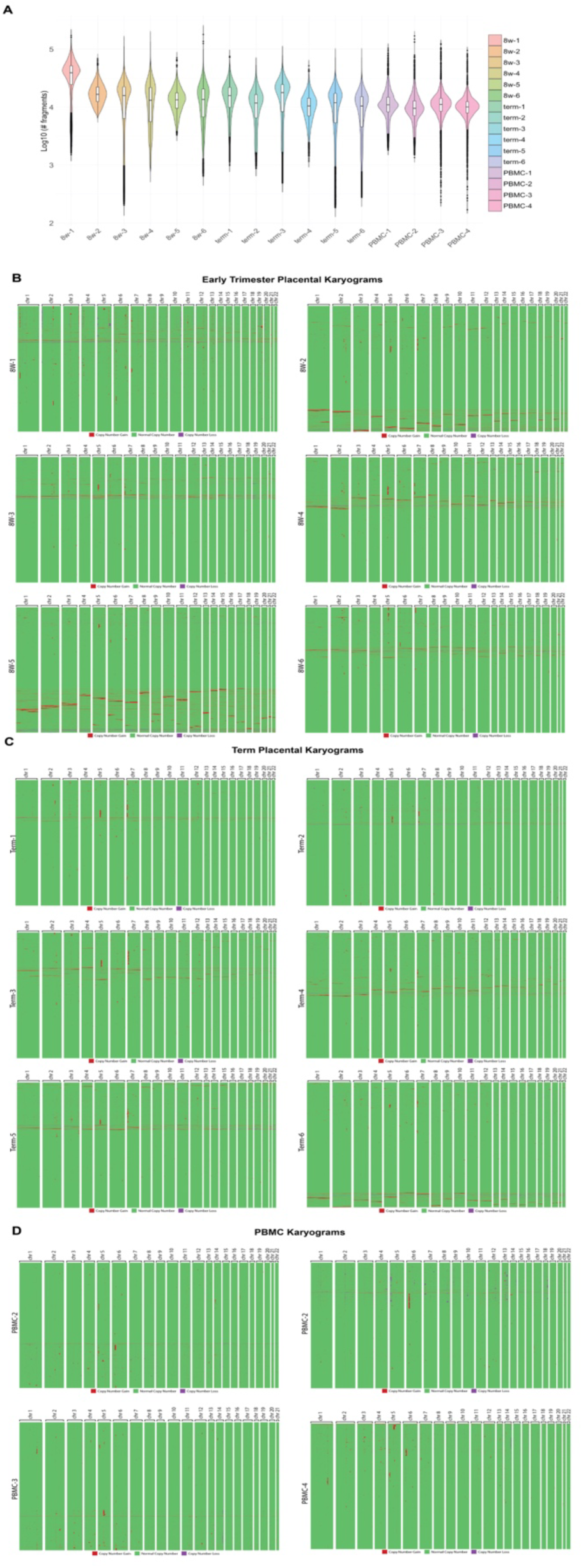
Detection of copy number alterations and aneuploidy in human placenta by snATAC (related to Figure 6.) (A) Violin plot of log-normalized fragment counts, illustrating the distribution of read counts across cells. (B) CNA karyograms for CT and ST nuclei from Tirst-trimester placental villi from six individual normal pregnancy. (C) CNA karyograms for CT and ST nuclei from term placental villi from six individual normal pregnancy. (D) CNA karyograms for human PBMCs from four healthy individuals.

## References

1. Del Gobbo, G. F. et al. Genomic imbalances in the placenta are associated with poor fetal growth. Molecular Medicine 27, 3 (2021).

2. Malvestiti, F. et al. Interpreting mosaicism in chorionic villi: results of a monocentric series of 1001 mosaics in chorionic villi with follow-up amniocentesis. Prenatal Diagnosis 35, 1117– 1127 (2015).

3. Hahnemann, J. M. & Vejerslev, L. O. European collaborative research on mosaicism in CVS (EUCROMIC)—fetal and extrafetal cell lineages in 192 gestations with CVS mosaicism involving single autosomal trisomy. American Journal of Medical Genetics 70, 179–187 (1997).

4. Pittalis, M. C. et al. The predictive value of cytogenetic diagnosis after CVS based on 4860 cases with both direct and culture methods. Prenatal Diagnosis 14, 267–278 (1994).

5. Ledbetter, D. H. et al. Cytogenetic results from the U.S. Collaborative Study on CVS. Prenat Diagn 12, 317–345 (1992).

6. Weier, J. F. et al. Human cytotrophoblasts acquire aneuploidies as they differentiate to an invasive phenotype. Developmental Biology 279, 420–432 (2005).

7. Weier, J. F. et al. Analysis of human invasive cytotrophoblasts demonstrates mosaic aneuploidy. PLOS ONE 18, e0284317 (2023).

8. Eggenhuizen, G. M., Go, A., Koster, M. P. H., Baart, E. B. & Galjaard, R. J. Confined placental mosaicism and the association with pregnancy outcome and fetal growth: a review of the literature. Human Reproduction Update 27, 885–903 (2021).

9. Spinillo, S. L. et al. Pregnancy outcome of confined placental mosaicism: meta-analysis of cohort studies. American Journal of Obstetrics and Gynecology 227, 714–727.e1 (2022).

10. Okae, H. et al. Derivation of Human Trophoblast Stem Cells. Cell Stem Cell 22, 50–63.e6 (2018).

11. Bai, T. et al. Establishment of human induced trophoblast stem-like cells from term villous cytotrophoblasts. Stem Cell Research 56, 102507 (2021).

12. Horii, M., Bui, T., Touma, O., Cho, H. Y. & Parast, M. M. An Improved Two-Step Protocol for Trophoblast Differentiation of Human Pluripotent Stem Cells. Curr Protoc Stem Cell Biol 50, e96 (2019).

13. Dong, C. et al. Derivation of trophoblast stem cells from naïve human pluripotent stem cells. eLife 9, e52504 (2020).

14. Cinkornpumin, J. K. et al. Naive Human Embryonic Stem Cells Can Give Rise to Cells with a Trophoblast-like Transcriptome and Methylome. Stem Cell Reports 15, 198–213 (2020).

15. Castel, G. et al. Induction of Human Trophoblast Stem Cells from Somatic Cells and Pluripotent Stem Cells. Cell Reports 33, 108419 (2020).

16. Sheridan, M. A. et al. Characterization of primary models of human trophoblast. Development 148, dev199749 (2021).

17. Guo, G. et al. Human naive epiblast cells possess unrestricted lineage potential. Cell Stem Cell 28, 1040–1056.e6 (2021).

18. Io, S. et al. Capturing human trophoblast development with naive pluripotent stem cells in vitro. Cell Stem Cell 28, 1023–1039.e13 (2021).

19. Wei, Y. et al. Efficient derivation of human trophoblast stem cells from primed pluripotent stem cells. Science Advances 7, eabf4416 (2021).

20. Jang, Y. J., Kim, M., Lee, B.-K. & Kim, J. Induction of human trophoblast stem-like cells from primed pluripotent stem cells. Proceedings of the National Academy of Sciences 119, e2115709119 (2022).

21. Kobayashi, N. et al. The microRNA cluster C19MC confers differentiation potential into trophoblast lineages upon human pluripotent stem cells. Nat Commun 13, 3071 (2022).

22. Soncin, F. et al. Derivation of functional trophoblast stem cells from primed human pluripotent stem cells. Stem Cell Reports 17, 1303–1317 (2022).

23. Viukov, S. et al. Human primed and naïve PSCs are both able to differentiate into trophoblast stem cells. Stem Cell Reports 17, 2484–2500 (2022).

24. Karvas, R. M., David, L. & Theunissen, T. W. Accessing the human trophoblast stem cell state from pluripotent and somatic cells. Cell. Mol. Life Sci. 79, 604 (2022).

25. Zorzan, I. et al. Chemical conversion of human conventional PSCs to TSCs following transient naive gene activation. EMBO reports **n/a**, e55235 (2023).

26. De Paepe, C. et al. BMP4 plays a role in apoptosis during human preimplantation development. Mol Reprod Dev 86, 53–62 (2019).

27. Guo, G. et al. Epigenetic resetting of human pluripotency. Development 144, 2748–2763 (2017).

28. Guo, G. et al. Human naive epiblast cells possess unrestricted lineage potential. Cell Stem Cell 28, 1040–1056.e6 (2021).

29. Yu, L. et al. Blastocyst-like structures generated from human pluripotent stem cells. Nature 591, 620–626 (2021).

30. Kagawa, H. et al. Human blastoids model blastocyst development and implantation. Nature 601, 600–605 (2022).

31. Roberts, R. M. et al. The role of BMP4 signaling in trophoblast emergence from pluripotency. Cellular and Molecular Life Sciences: CMLS 79, 447 (2022).

32. Zhang, P. et al. Short-term BMP-4 treatment initiates mesoderm induction in human embryonic stem cells. Blood 111, 1933–1941 (2008).

33. Kurek, D. et al. Endogenous WNT signals mediate BMP-induced and spontaneous differentiation of epiblast stem cells and human embryonic stem cells. Stem Cell Reports 4, 114–128 (2015).

34. Martyn, I., Kanno, T. Y., Ruzo, A., Siggia, E. D. & Brivanlou, A. H. Self-organization of a human organizer by combined Wnt and Nodal signalling. Nature 558, 132–135 (2018).

35. Amita, M. et al. Complete and unidirectional conversion of human embryonic stem cells to trophoblast by BMP4. Proceedings of the National Academy of Sciences of the United States of America 110, E1212 (2013).

36. Yang, Y. et al. Heightened potency of human pluripotent stem cell lines created by transient BMP4 exposure. Proceedings of the National Academy of Sciences of the United States of America 112, E2337 (2015).

37. Lannoo, L. et al. Rare autosomal trisomies detected by non-invasive prenatal testing: an overview of current knowledge. Eur J Hum Genet 1–8 (2022) doi:10.1038/s41431-022-01147-1.

38. Yang, M. et al. Evaluation of genome-wide DNA methylation profile of human embryos with different developmental competences. Human Reproduction 36, 1682–1690 (2021).

39. Qi, Y. et al. The significance of trisomy 7 mosaicism in noninvasive prenatal screening. Human Genomics 13, 18 (2019).

40. Zhu, X. et al. Clinical Significance of Non-Invasive Prenatal Screening for Trisomy 7: Cohort Study and Literature Review. Genes 12, 11 (2020).

41. Thompson, S. L. & Compton, D. A. Proliferation of aneuploid human cells is limited by a p53-dependent mechanism. The Journal of Cell Biology 188, 369 (2010).

42. Bakhoum, S. F. & Cantley, L. C. The multifaceted role of chromosomal instability in cancer and its microenvironment. Cell 174, 1347–1360 (2018).

43. Barrio, L. et al. Chromosomal instability-induced cell invasion through caspase-driven DNA damage. Current Biology 33, 4446–4457.e5 (2023).

44. Burrell, R. A. et al. Replication stress links structural and numerical cancer chromosomal instability. Nature 494, 492–496 (2013).

45. Passerini, V. et al. The presence of extra chromosomes leads to genomic instability. Nat Commun 7, 10754 (2016).

46. Karvas, R. M. et al. Stem-cell-derived trophoblast organoids model human placental development and susceptibility to emerging pathogens. Cell Stem Cell 29, 810–825.e8 (2022).

47. Shannon, M. J. et al. Single-cell assessment of primary and stem cell-derived human trophoblast organoids as placenta-modeling platforms. Developmental Cell 59, 776–792.e11 (2024).

48. Turco, M. Y. et al. Trophoblast organoids as a model for maternal–fetal interactions during human placentation. Nature 564, 263–267 (2018).

49. Sheridan, M. A. et al. Establishment and differentiation of long-term trophoblast organoid cultures from the human placenta. Nat Protoc 15, 3441–3463 (2020).

50. Schroeder, D. I. et al. The human placenta methylome. Proceedings of the National Academy of Sciences 110, 6037–6042 (2013).

51. Hemberger, M., Udayashankar, R., Tesar, P., Moore, H. & Burton, G. J. ELF5-enforced transcriptional networks define an epigenetically regulated trophoblast stem cell compartment in the human placenta. Hum Mol Genet 19, 2456–2467 (2010).

52. Weissbein, U., Benvenisty, N. & Ben-David, U. Genome maintenance in pluripotent stem cells. Journal of Cell Biology 204, 153–163 (2014).

53. Ben-David, U., Benvenisty, N. & Mayshar, Y. Genetic instability in human induced pluripotent stem cells: Classification of causes and possible safeguards. Cell Cycle 9, 4603–4604 (2010).

54. Wu, J. & Barbaric, I. Fitness selection in human pluripotent stem cells and interspecies chimeras: Implications for human development and regenerative medicine. Developmental Biology 476, 209 (2021).

55. Leylek, T. R., Jeusset, L. M., Lichtensztejn, Z. & McManus, K. J. Reduced Expression of Genes Regulating Cohesion Induces Chromosome Instability that May Promote Cancer and Impact Patient Outcomes. Sci Rep 10, 592 (2020).

56. Michel, L. S. et al. MAD2 haplo-insufficiency causes premature anaphase and chromosome instability in mammalian cells. Nature 409, 355–359 (2001).

57. Wang, X. et al. Correlation of defective mitotic checkpoint with aberrantly reduced expression of MAD2 protein in nasopharyngeal carcinoma cells. Carcinogenesis 21, 2293– 2297 (2000).

58. Dobles, M., Liberal, V., Scott, M. L., Benezra, R. & Sorger, P. K. Chromosome Missegregation and Apoptosis in Mice Lacking the Mitotic Checkpoint Protein Mad2. Cell 101, 635–645 (2000).

59. Xu, H. et al. Rad21-Cohesin Haploinsufficiency Impedes DNA Repair and Enhances Gastrointestinal Radiosensitivity in Mice. PLOS ONE 5, e12112 (2010).

60. Xu, H. et al. Enhanced RAD21 cohesin expression confers poor prognosis and resistance to chemotherapy in high grade luminal, basal and HER2 breast cancers. Breast Cancer Research 13, R9 (2011).

61. Levy, R. et al. Trophoblast apoptosis from pregnancies complicated by fetal growth restriction is associated with enhanced p53 expression. American Journal of Obstetrics and Gynecology 186, 1056–1061 (2002).

62. Sherr, C. J. & Roberts, J. M. CDK inhibitors: positive and negative regulators of G1-phase progression. Genes Dev. 13, 1501–1512 (1999).

63. Blain, S. Switching cyclin D-Cdk4 kinase activity on and off. Cell Cycle 7, 892–898 (2008).

64. Serrano, M. The INK4a/ARF locus in murine tumorigenesis. Carcinogenesis 21, 865–869 (2000).

65. Beishline, K. & Azizkhan-Clifford, J. Sp1 and the ‘hallmarks of cancer’. The FEBS Journal 282, 224–258 (2015).

66. Liu, D. et al. Autophagy regulates the survival of cells with chromosomal instability. Oncotarget 7, 63913–63923 (2016).

67. Yun, C. W., Jeon, J., Go, G., Lee, J. H. & Lee, S. H. The Dual Role of Autophagy in Cancer Development and a Therapeutic Strategy for Cancer by Targeting Autophagy. International Journal of Molecular Sciences 22, 179 (2020).

68. Singla, S., Iwamoto-Stohl, L. K., Zhu, M. & Zernicka-Goetz, M. Autophagy-mediated apoptosis eliminates aneuploid cells in a mouse model of chromosome mosaicism. Nat Commun 11, 2958 (2020).

69. Tang, Y.-C., Williams, B. R., Siegel, J. J. & Amon, A. The energy and proteotoxic stress-inducing compounds AICAR and 17-AAG antagonize proliferation in aneuploid cells. Cell 144, 499 (2011).

70. Liu, Y. et al. Single-cell RNA-seq reveals the diversity of trophoblast subtypes and patterns of differentiation in the human placenta. Cell Res 28, 819–832 (2018).

71. Wang, M. et al. Single-nucleus multi-omic profiling of human placental syncytiotrophoblasts identifies cellular trajectories during pregnancy. Nat Genet 56, 294–305 (2024).

72. Griffiths, J. A., Scialdone, A. & Marioni, J. C. Mosaic autosomal aneuploidies are detectable from single-cell RNAseq data. BMC Genomics 18, 904 (2017).

73. Yang, M. et al. Depletion of aneuploid cells in human embryos and gastruloids. Nature Cell Biology 23, 15 (2021).

74. Nikolic, A. et al. Copy-scAT: Deconvoluting single-cell chromatin accessibility of genetic subclones in cancer. Science Advances 7, eabg6045 (2021).

75. Moore, T. W. & Yardımcı, G. G. Robust CNV detection using single-cell ATAC-seq. Preprint at 10.1101/2023.10.04.560975 (2023).

76. Ramakrishnan, A. et al. epiAneufinder identifies copy number alterations from single-cell ATAC-seq data. Nat Commun 14, 5846 (2023).

77. Coorens, T. H. H. et al. Inherent mosaicism and extensive mutation of human placentas. Nature 592, 80–85 (2021).

78. Wallace, A. D. et al. Placental somatic mutation in human stillbirth and live birth: A pilot case-control study of paired placental, fetal, and maternal whole genomes. Placenta 154, 137–144 (2024).

79. Thompson, S. L., Bakhoum, S. F. & Compton, D. A. Mechanisms of Chromosomal Instability. Current biology : CB 20, R285 (2010).

80. Hosea, R., Hillary, S., Naqvi, S., Wu, S. & Kasim, V. The two sides of chromosomal instability: drivers and brakes in cancer. Signal Transduction and Targeted Therapy 9, 1– 30 (2024).

81. Zhou, P., Wang, J., Wang, J. & Liu, X. When autophagy meets placenta development and pregnancy complications. Front. Cell Dev. Biol. 12, (2024).

82. Cecconi, F. & Levine, B. The Role of Autophagy in Mammalian Development: Cell Makeover Rather than Cell Death. Developmental cell 15, 344 (2008).

83. Deglincerti, A. et al. Self-organization of human embryonic stem cells on micropatterns. Nat Protoc 11, 2223–2232 (2016).

84. De Santis, R. et al. The emergence of human gastrulation upon in vitro attachment. Stem Cell Reports 19, 41–53 (2024).

85. Cock, P. J. A., Fields, C. J., Goto, N., Heuer, M. L. & Rice, P. M. The Sanger FASTQ file format for sequences with quality scores, and the Solexa/Illumina FASTQ variants. Nucleic Acids Research 38, 1767–1771 (2010).

86. Langmead, B. & Salzberg, S. L. Fast gapped-read alignment with Bowtie 2. Nat Methods 9, 357–359 (2012).

87. Li, B. & Dewey, C. N. RSEM: accurate transcript quantification from RNA-Seq data with or without a reference genome. BMC Bioinformatics 12, 323 (2011).

88. Love, M. I., Huber, W. & Anders, S. Moderated estimation of fold change and dispersion for RNA-seq data with DESeq2. Genome Biol 15, 550 (2014).

89. Cock, P. J. A., Fields, C. J., Goto, N., Heuer, M. L. & Rice, P. M. The Sanger FASTQ file format for sequences with quality scores, and the Solexa/Illumina FASTQ variants. Nucleic Acids Research 38, 1767–1771 (2010).

90. Krueger, F. & Andrews, S. R. Bismark: a flexible aligner and methylation caller for Bisulfite-Seq applications. Bioinformatics 27, 1571–1572 (2011).

91. Lister, R. et al. Human DNA methylomes at base resolution show widespread epigenomic differences. Nature 462, 315–322 (2009).

92. Okae, H. et al. Derivation of Human Trophoblast Stem Cells. Cell Stem Cell 22, 50–63.e6 (2018).

93. R Core Team. R: A Language and Environment for Statistical Computing. R Foundation for Statistical Computing (2024).

94. Wickham, H. ggplot2. WIREs Computational Statistics 3, 180–185 (2011).

95. Bates, D., Mächler, M., Bolker, B. & Walker, S. Fitting Linear Mixed-Effects Models using lme4. Preprint at 10.48550/arXiv.1406.5823 (2014).

96. Arel-Bundock, V., Greifer, N. & Heiss, A. How to Interpret Statistical Models Using marginaleffects in R and Python. Journal of Statistical Software ((Forthcoming)).

97. Ahlmann-Eltze, C. & Patil, I. ggsignif: R Package for Displaying Significance Brackets for ‘ggplot2’. Preprint at 10.31234/osf.io/7awm6 (2021).

98. Patro, R., Duggal, G., Love, M. I., Irizarry, R. A. & Kingsford, C. Salmon provides fast and bias-aware quantification of transcript expression. Nat Methods 14, 417–419 (2017).

99. Soneson, C., Love, M. I. & Robinson, M. D. Differential analyses for RNA-seq: transcript-level estimates improve gene-level inferences. Preprint at 10.12688/f1000research.7563.1 (2016).

100. Hao, Y. et al. Dictionary learning for integrative, multimodal and scalable single-cell analysis. Nat Biotechnol 42, 293–304 (2024).

