## Supplementary material for "Chromosomal Instability in Human Trophoblast Stem Cells and Placentas": Table S1

| Cell line | G-banding metaphase cells |
|---|---|
| nTSC P11 | 46 XX [15]* |
| | 43 XX, -4, -9, -18 [1] |
| | 44 XX, -18, -19, -22, +mar [1] |
| | 45 XX, -1 [1] |
| | 45 XX, -2 [1] |
| | 45 XX, -22 [1] |
| nbTSC P13 #1 | 46 XX [15] 15 normal female karyotype |
| | 43 XX, -9, -11,-17 [1] |
| | 43 XX, -3. -4,-15, del(17) (p10) [1] |
| | 45 XX, -17, del (20) (q10) [1] |
| | 45 XX, -13 [1] |
| | 46 XX, -17, +19, del (20)(p11.2)[1] |
| nbTSC P13 #2 | A female karyotype with chromosome counts from 81-95 [3] |
| ccTSC p12 #1 | 47 XX, +mar [15] |
| | 46 X, -X, +mar [2] |
| | 46 XX, -22, +mar [1] |
| | 48 XX, +18, +mar [1] |
| | 48 XX, +16, +mar [1] |
| ccTSC P10 #2 | 47 XX, +7 [6] |
| | 48 XX, +7, +7 [2] |
| | 46 XX, +7, +12, -14, -18 [1] |
| | 48 XX, +2 +7, add(15)(q24)….[1] |
| | 47XX, -1 +7 +11[1] |
| | 48 XX, +7, +11, del(18)(p10)[1] |
| | 48XX +2 +7 [1] |
| | 47 XX +7,+12, -18[1] |
| | 49 XX, -5, +7, +11, +18[1] |
| | 52 XX, +3 +5 +6 +7, +13, +15, +18, -21[1] |
| | 48 XX +7 +20[1] |
| | 43 XX, +7, -8, -12, -19, -21[1] |

| | |
|---|---|
| | 52 XX, -1, +2, +3, +7, +7, +11,+14,+16, -19, +22[1] |
| | 49XX, +X, +5, +7, -10, +11[1] |
| pdTSC P12 | 46 XX [10] 10 normal female karyotype |
| | 46 XX, t(1, 13)(q32.3; q12.1) [9] |
| | 46 XX, t(1, 13)(q32.3; q12.1), add (11) (p11.2) [1] |

* There is the cell number within the bracket
