## Supplementary material for "Chromosomal Instability in Human Trophoblast Stem Cells and Placentas": Table S2

| Cluster | Rate (%) | # of cells | Residuals | nGenes | Zeros |
|---|---|---|---|---|---|
| EVT8W_1 | 9.6 | 52 | 0.282 | 4190 | Poor |
| EVT8W_2 | 6.7 | 193 | 0.355 | 3756 | Poor |
| EVT8W_3 | 15 | 280 | 0.374 | 3137 | Poor |
| EVT24W | 9.2 | 195 | 0.203 | 3016 | Good |
| CTB_1 | 27.7 | 159 | 0.367 | 1907 | Poor |
| CTB_2 | 37.1 | 70 | 0.57 | 1077 | Poor |
| CTB_3 | 3.3 | 61 | 0.375 | 3121 | Poor |
| Min.median = 19 | | | | | |
