## Supplementary material for "Chromosomal Instability in Human Trophoblast Stem Cells and Placentas": Table S3

**Table S3. Antibodies**

| | | |
|---|---|---|
| Mouse monoclonal TFAP2A | DSHB | 3B5S |
| Mouse monoclonal P62 | R&D | MAB8028 |
| Mouse monoclonal LAMP1 | R&D | MAB4800 |
| Mouse monoclonal TFAP2C | Santa Cruz | sc-12762 |
| Mouse monoclonal GATA4 | Santa Cruz | sc-25310 |
| Mouse Polyclonal GATA3 | Invitrogen | MA1-028 |
| Rabbit monoclonal KRT7 | Abcam | ab183344 |
| Rabbit monoclonal HLA-G | Cell-sig | 79769 |
| Rabbit polyclonal Beclin1 | Novus | NB500-249SS |
| Rabbit monoclonal LC3B | R&D | MAB85582 |
| Rabbit monoclonal GATA6 | Cell-sig | D61E4 |
| Rabbit monoclonal CDX2 | Cell-sig | D11D10 |
| Rabbit polyclonal KLF17 | Sigma | HPA024629 |
| Rabbit polyclonal yH2AX | Novus Biologicals | NB100-384 |
| Goat polyclonal SDC1 | R&D | AF2780 |
| Goat polyclonal Cathepsin B | R&D | AF953 |
| Rabbit polyclonal B-Actin | Proteintech | 20536-1-AP |
| Goat polyclonal GATA2 | R&D | AF2046 |
| Goat polyclonal OCT3/4 | R&D | AF1759 |
| Goat polyclonal SOX17 | R&D | AF1924 |
| Goat Polyclonal NANOG | R&D | AF1997 |
| Donkey anti-mouse Alexa 647 | Invitrogen | A31571 |
| Donkey anti-rabbit Alexa 488 | Invitrogen | A21206 |
| Donkey anti-goat Alexa 594 | Invitrogen | A11058 |
| Donkey anti-goat Alexa 555 | Invitrogen | A11058 |
| Mouse IgG HRP-conjugated Antibo | R&D Systems | HAF007 |
| Rabbit IgG HRP-conjugated Antibo | R&D Systems | HAF008 |
| Goat IgG HRP-conjugated Antibody | R&D Systems | HAF109 |
