## Supplementary material for "Chromosomal Instability in Human Trophoblast Stem Cells and Placentas": Table S4

| Gene | Forward | Reverse |
|---|---|---|
| NANOG | GCAGAAGGCCTCAGCACCTA | AGGTTCCCAGTCGGGTTCA |
| POU5F1/OCT4 | GCTCGAGAAGGATGTGGTCC | CGTTGTGCATAGTCGCTGCT |
| CGB3 | ACCCTGGCTGTGGAGAAGGAG | ATGGACTCGAAGCGCACATCG |
| SDC1 | CTTCACACTCCCCACACAGA | GTATTCTCCCCCGAGGTTTC |
| PSG1 | GGTACAAAGGGCAAATGAGG | ATTCTGGATCAGCAGGGATG |
| INHA1 | ATCCTTTTCCCAGCCACAG | GCCGGAACATGTATCTGAAG |
| GATA2 | GACTACAGCAGCGGACTCTT | GCCTTCTGAACAGGAACGAG |
| PTGES | AGGATGCCCTGAGACACGGA | CCAGGAAAAGGAAGGGGTAG |
| TP63 | ACGAAGATCCCCAGATGATG | TGCTGTTGCCTGTACGTTTC |
| KRT19 | ACCTGGAGATGCAGATCGAA | AATCCACCTCCACACTGACC |
| VIM | AGTCCACTGAGTACCGGAGAC | CATTTCACGCATCTGGCGTTC |
| CHD10 | AGATGCCGATGACCCTTCATA | TGTTCGGTAAAGCAGTCCTGA |
| IGFBP3 | CCTGCCGTAGAGAAATGGAA | AAGGGCGACACTGCTTTTT |
| VTCN1 | TCTGGGCATCCCAAGTTGAC | TCCGCCTTTTGATCTCCGATT |
| HLA-G | GAGGAGACACGGAACACCAAG | GTCGCAGCCAATCATCCACT |
| NOTUM | TTTGGCTACAAGGTCTACCCG | TCAAACAGCCACTGCACCAC |
| LRRC32 | GCTGCACAACACCAAGACAAA | GATCAAGGGTCTCAGTGTCTGG |
| CSH1/2 | ATTTCTGTTGCGTTTCCTCCAT | CATGACTCCCAGACCTCCTTCT |
| CGA | CACTCCACTAAGGTCCAAGAAGA | CCGTGTGGTTCTCCACTTTGA |
| SDC1 | CTTCACACTCCCCACACAGA | GTATTCTCCCCCGAGGTTTC |
| CGB | ACCCTGGCTGTGGAGAAGGAG | ATGGACTCGAAGCGCACATCG |
| PSG3 | TCGTAAAGCGAGGTGATGGG | AAGCTCACAGCCTCCATGTC |
| KRT7 | AGGATGTGGATGCTGCCTAC | CACCACAGATGTGTCGGAGA |
| GATA3 | CTACTACGGAAACTCGGTCAGGGC | AGCCAGGGTAGGGATCCATGAAG |
| TFAP2C | ACGAAATGAGATGGCAGCTAGGAAGA | TGTCCGGTCTTGGCTGAGAAGTTC |
| GAPDH | AGGGCTGCTTTTAACTCTGGT | CCCCACTTGATTTTGGAGGGA |
